# OptoPi: An open source flexible platform for the analysis of small animal behaviour

**DOI:** 10.1101/2022.07.12.499786

**Authors:** Xavier Cano-Ferrer, Ruairí J.V. Roberts, Alice S. French, Joost de Folter, Hui Gong, Luke Nightingale, Amy Strange, Albane Imbert, Lucia L. Prieto-Godino

## Abstract

Behaviour is the ultimate output of neural circuit computations, and therefore its analysis is a cornerstone of neuroscience research. However, every animal and experimental paradigm requires different illumination conditions to capture and, in some cases, manipulate specific behavioural features. This means that researchers often develop, from scratch, their own solutions and experimental set-ups. Here, we present OptoPi, an open source, affordable (∼ £600), behavioural arena with accompanying multi-animal tracking software. The system features highly customisable and reproducible visible and infrared illumination and allows for temporally precise optogenetic stimulation. OptoPi acquires images using a Raspberry Pi camera, features motorised LED-based illumination, Arduino control, as well as spectrum and irradiance monitoring to fine-tune illumination conditions with real time feedback. Our open-source software (BIO) can be used to simultaneously track multiple animals while accurately keeping individual animal’s identity both in on-line and off-line modes. We demonstrate the functionality of OptoPi by recording and tracking under different illumination conditions the spontaneous behaviour of larval zebrafish as well as adult *Drosophila* flies and their first instar larvae, an experimental animal that due to its small size and transparency has classically been hard to track. Further, we showcase OptoPi’s optogenetic capabilities through a series of experiments using transgenic *Drosophila* larvae.

## 1. Hardware in context

A key method in neuroscience is behavioural analysis, including the manipulation of neuronal activity while monitoring behavioural output. A commonly used approach to accurately modify neuronal activity is optogenetics, whereby animals are genetically engineered to express light-sensitive ion channels in specific neuronal populations. This method is particularly accessible for small animals such as *Drosophila* and zebrafish where stimulation can be delivered in freely behaving individuals.

Over the last decade numerous open hardware solutions have been published to facilitate the study of small animal behaviour. These range from relatively generic and modular systems such as the FlyPi [1], able to record and optogenetically stimulate animals, to more sophisticated and task-specific set-ups. For example, Ethoscopes were designed specifically to measure and manipulate the sleep of flies in real time through close-loop experiments, but can also be adapted for other assays [2]. Another excellent example is PiVR the first open source behavioural device designed to perform closed loop experiments with light stimulation [3]. However, a limitation of these previous designs is that although their performance is excellent for the specific task for which they were developed, adapting them to different purposes, such as using different experimental animals, often requires modifications of the illumination and/or stimulation conditions, some of which can be difficult or time-consuming to implement.

To overcome these limitations, here we present a hardware solution that enables the experimenter to change the illumination, recording, and optogenetic stimulation conditions flexibly and rapidly. Our platform consists of a stage to place the animal arena and multiple movable illumination modules whose angle, light intensity and wavelength can be adjusted in a matter of seconds with user-friendly knobs. Additionally, the camera distance to the arena can also be easily modified. Together these adaptations enable rapid control over recording conditions to find settings that are optimal for each experiment. Our device is also fitted with multiple wavelength LEDs that match the activation spectrum of the most popularly used optogenetic actuators. These are placed onto adjustable modules to ensure optimal optogenetic stimulation. To quickly reproduce previous recording and stimulation settings, OptoPi is equipped with spectrum, irradiance and ultrasound sensors that monitor illumination conditions and placement of illumination modules. Finally, we also describe the development of a new open-source tracking software, which can be used in on and off-line modes, and that enables the tracking of multiple animals while keeping individual identity after collisions.

We anticipate that the flexibility of our platform will enable researchers to have a single solution for multiple behavioural analysis needs within their lab and will help the standardisation of behavioural assays in small animals. In addition, OptoPi’s hardware could be combined with previously published modules to expand its functionality. For example, the camera recording settings could be controlled using the graphical using interface (GUI) of the FlyPi, or our hardware could be combined with the software capabilities of PiVR for close-loop experiments. Therefore, OptoPi expands the open source toolkit that neuroscience researchers can take advantage of to perform behavioural analyses in a wide variety of animals and settings.

## 2. Hardware description

This section has the following subsections:

2.1. Overview
2.2. Illumination control
2.3 Optogenetic modules
2.4 Recording settings
2.5 Tracking options

### 2.1. Overview

OptoPi consists of a mechanical framework, composed of 3D printed, laser-cut and mechanical parts such as the four metal posts (Figure 1a). The behavioural arena is placed on a static platform, that can have its height adjusted, while the illumination, stimulation and camera modules are movable to achieve optimal experimental conditions. The device is controlled by an Arduino via manual knobs for maximum ease of use (Figure 1b). Videos or images are captured via a Raspberry Pi Camera and can be processed through several programs. However, here we present the development of BIO, an open source software that can perform tracking of multiple individuals and resolve identities after collisions.

**Figure 1.**
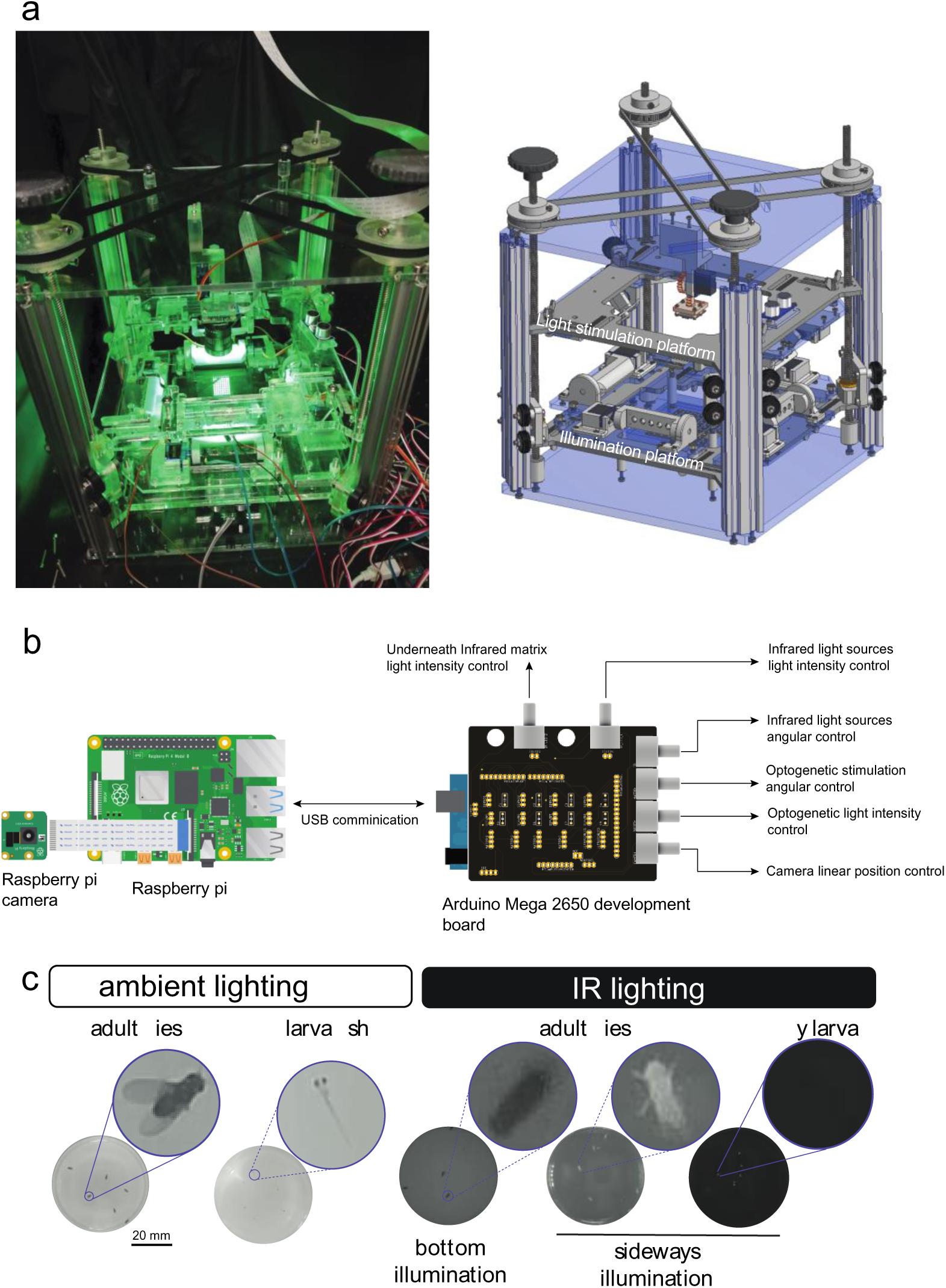
Fly arena design. **a.** Mechanical assembly of OptoPi using CAD software on the right. The stimulation and illumination platforms that surround the arena move freely and independently. Fully assembled arena on the left with the Raspberry Pi HQ camera visible in the centre. **b.** The Raspberry Pi both controls the camera, which records the experiment, and communicates via serial port with the Arduino Mega 2560. The Arduino Mega 2560 has its own shielding to allow the researcher to control the source, position and intensity of light as well as the camera position. **c.** Examples of different lighting configurations. Ambient lighting works best for dark animals such as adult flies and larvae zebrafish when no optogenetic manipulation is required. For optogenetic manipulation and transparent animals such as fly larvae, IR lighting is preferred.

### 2.2. Illumination control

The arena is a laser cut Polymethyl methacrylate (PMMA) rectangular platform (120 x 120 mm) that can be located at different heights by using 3D printed spacers. This makes it possible to use different materials to change the diffusion of backlit infrared (IR) or white illumination. We provide a few designs that create different illumination or stimulation conditions by positioning IR and RGB LEDs at different levels (Figure 1c). The dimensions of the arena provide enough surface to image small animals in a wide range of platforms such as petri dishes 60-100 mm or custom arenas such as the ones proposed by Koemans [4] or Gou [5].

The illumination modules have been designed to enable control of the angular and linear (horizontal and vertical) position, as well as light intensity of all the light sources: Infrared for illumination of experiments in the dark, for example when performing optogenetics, and RGB for white light illumination and optogenetic stimulation. The device has a I^2^C port to connect an illuminance sensor that enables the user to characterize and adapt it to new light stimulation tools such as other RGB matrices, or LED displays. In addition, a spectral sensor can be connected but we did not use it in this manuscript because we found the resolution not useful for the experiments we were running.

The IR illumination is composed of four linear arrays surrounding the arena forming a square (hereafter ‘reflected IR modules’) and one matrix beneath the arena (hereafter ‘transmitted IR module’). Each linear array of the reflected IR modules is composed of five Infrared LED (AN5307B Stanley Electric, 940 nm IR LED, 5mm) and five current limiting resistors (68 Ω, 1% tolerance), that provide the nominal forward current of I_F_≈50 mA, soldered on a custom PCB provided in the publication files. Each array is mechanically connected to a micro servo motor, SG90, that enables angle control within a range of approximately 2π/3 radians (120°) with 1° of resolution by rotating a 10 kΩ potentiometer located on the right side of the control board (Figure 2a) ([Supplementary video 1](https://osf.io/ve976/)). The light intensity of the IR light sources can be controlled by an external 100 Ω, 5W potentiometer. Each light source is mounted on a guide that provides manual translation adjustment (20 mm) by tweaking the tightness of two M6 wing nuts. A scale in millimetres is engraved next to both guides (Figure 2b). The four light sources are held by a bigger platform that can move upwards and downwards (vertical translation movement) and therefore the height of the incident light can be changed.

**Figure 2.**
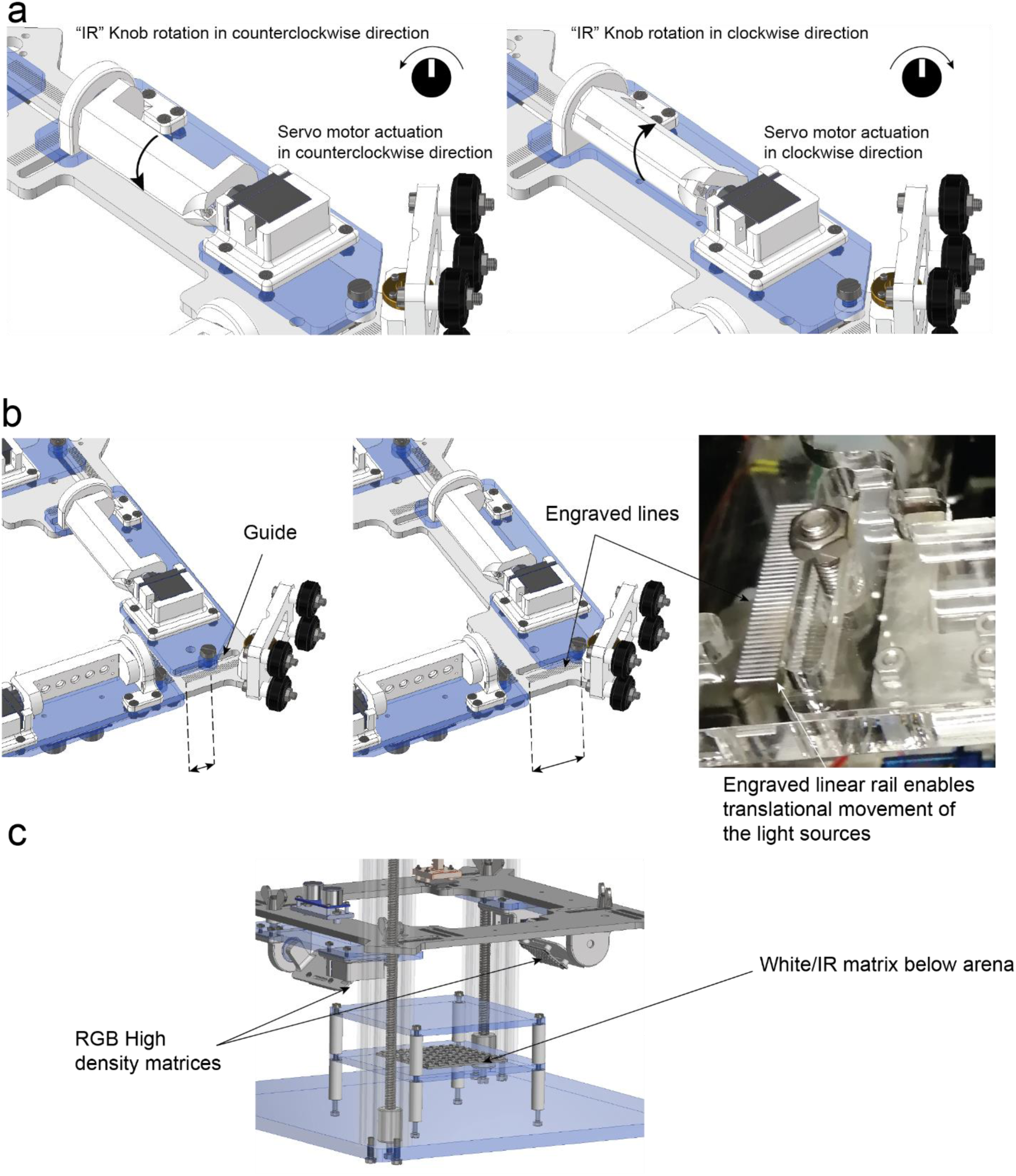
Movements that allow the customization of the device. **a.** Rotational movement of the Infrared light sources are adjusted by rotating the potentiometer analogue signal. **b**. The distance from all the light sources (RGB and IR) can be manually adjusted by untightening the M6 screws and sliding the light source platform forwards and backwards. **c.** Position of the RGB orientable matrices on the top platform and the arena underneath illumination.

An IR matrix is located beneath the arena to achieve backlight illumination when needed. It also has its own 100 Ω, 5W potentiometer to adjust light intensity by limiting the current. Additionally, this matrix can be easily interchanged with a white LED array in order to achieve white light transmitted illumination (Figure 2c).

Overhead white light illumination can be achieved by employing the “optogenetic” modules with all RGB LEDs on at the same level for each colour. This illumination is achieved by controlling two RGB Adafruit Dotstar high density 8×8 grids (Adafruit 3444). These grids are mounted on a second platform identical to the one used for the infrared illumination sources. Therefore, they can be adjusted to the same degree of freedom as the IR linear arrays angle, and are controlled by a separate RGB knob. Horizontal and vertical positions as well as their light intensity can also be changed with the corresponding potentiometer (Bright) (Figure 3a) ([Supplementary video 2](https://osf.io/jqh4z/)). The two platforms can be adjusted vertically by rotating a double GT2 timing belt-lead screw driven system (Figure 3b) ([Supplementary video 3](https://osf.io/cvswp/)). On each platform, an ultrasonic sensor HC - SR04 (Sparkfun 15569) measures that platform’s distance from the main structure: the illumination platform’s distance is relative to the bottom of the OptoPi and the stimulation platform relative to the top (Figure 3c). The signal received from the ultrasonic sensors is filtered with a moving average filter. This improves the signal to noise ratio and eases the interpretation of the readings by the user (Figure 3d).

**Figure 3.**
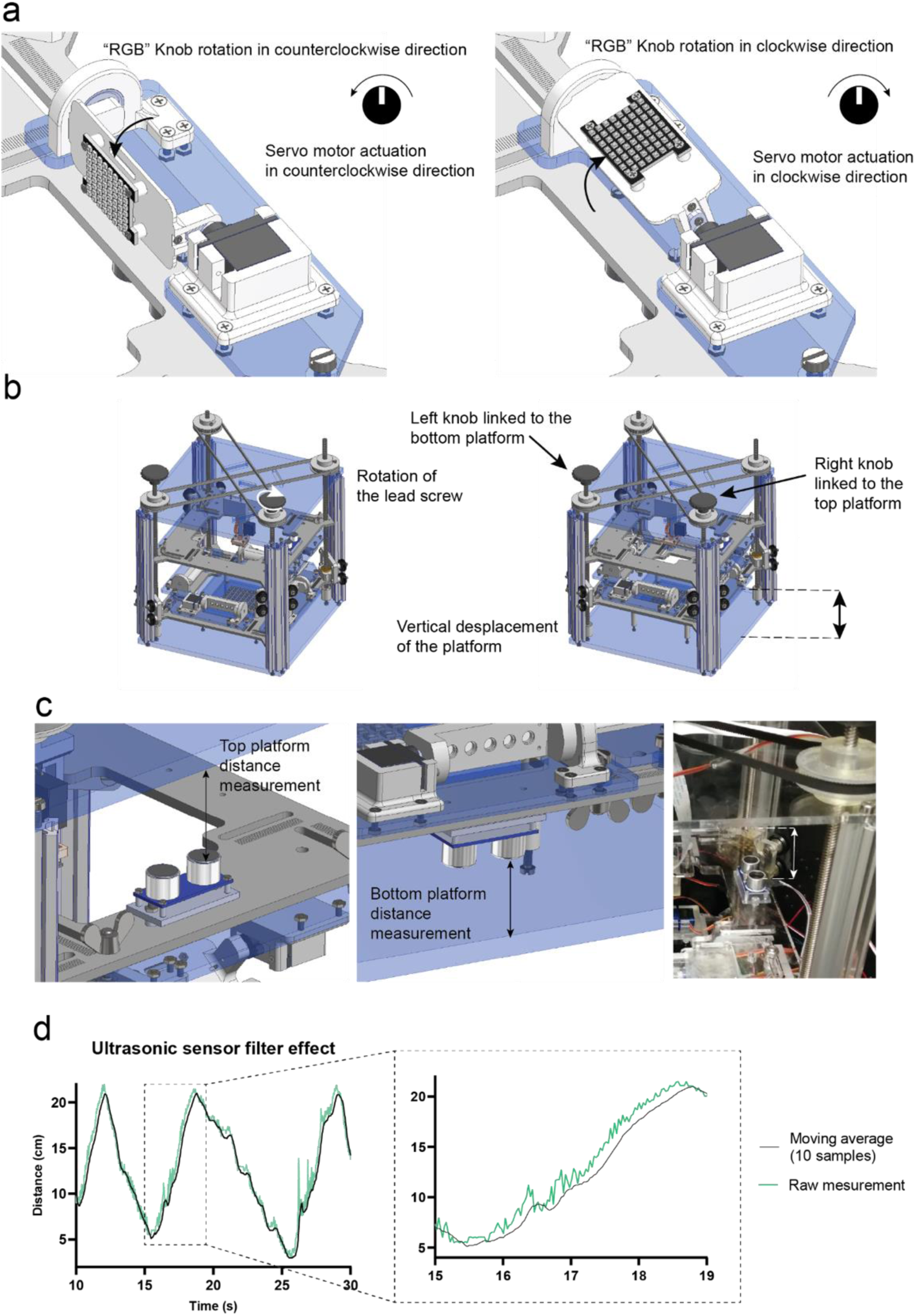
Customisation of light illumination. **a.** Rotational movement of the RGB matrices actuated by the analogue signal from the PCB potentiometer. **b.** The vertical position of the light source platforms (both IR and RGB) can be changed independently by rotating the two separate lead screws/ pulley systems. **c.** The ultrasonic sensors face the roof and the base of the OptoPi structure to measure the distance from each platform to its closest surface. These sensors determine the distance between the illumination platform and the OptoPi bottom and the distance between the stimulation platform and the OptoPi top. **d.** The moving average filter applied to the ultrasonic sensor signal results in less signal fluctuation making it easier for the user to read.

### 2.3 Optogenetic modules

The RGB illumination arrays can also be used as optogenetic modules. Optogenetics uses light to control the activity of cells that have been genetically modified to express light-sensitive ion channels. The optimal activation wavelengths of the most popularly used light-sensitive ion channels are 470, 490, 525 and 590 nm for GtACR2, ChR2^XXL^, GtACR1 and CsChrimson respectively [6]–[8]. In the present device we have measured the RGB matrices spectrum: 465,521 and 633 nm peak wavelengths using a Thorlabs CCS200/M compact spectrometer with the CCSB1 cosine corrector. These values fulfil the spectrum requirements for most of optogenetic experiments (Figure 4a).

**Figure 4.**
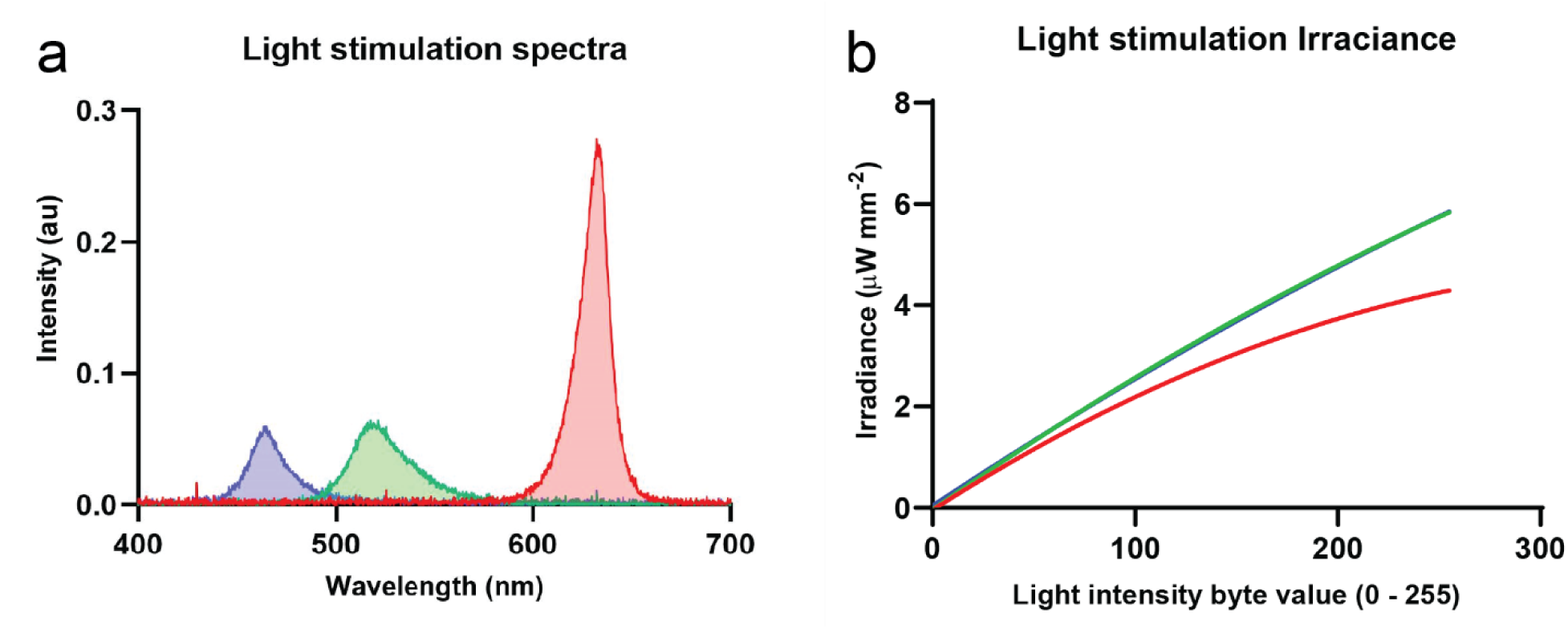
Optogenetic module characterization. **a.** Visible light spectrum of the RGB matrices measured with the Thorlabs CCS200/M compact spectrometer with the CCSB1 cosine corrector **b.** Irradiance measured on the setup using a platform with a calibrated Thorlabs S120VC photodiode connected to a PM100D compact power meter console.

Most recent optogenetic actuators require a light irradiance of 1 µW/mm^2^ or less [6]–[10] to be activated. Our device can generate maximum irradiances of 4.22, 5.88, 5.84 µW/mm^2^ for red, green and blue respectively. It does so by using the two RGB high density grids (Figure 4b). The OptoPi design leaves room for two more matrices which, can be added to boost the irradiance further if required. The measured average minimum change in irradiance is 0.024, 0.033, 0.033 µW/mm^2^ for the red, green, and blue LED channels respectively. These values have been measured using a calibrated Thorlabs S120VC photodiode and a PM100D power and energy meter. The measured residual irradiance (Brightness byte value 0) of the device is 0 1.5·10^-4^ µW/mm^2^.

Finally, the device has the capability to perform light intensity (TSL2561 Digital Luminosity Lux Light Sensor Module) measurements in the same platform arena where experimental animals will be placed. This enables precise measurement of the light stimulation intensity for any specific position or configuration of the RGB matrices. This feature, together with positional readings from the ultrasound sensors and motor angles of the LEDs, enables the precise recording of stimulation conditions. This includes the irradiance of the behavioural arena, allowing the quick reproduction of experimental settings within and across labs.

The main control shield is a custom PCB which has all the connections and required elements to drive the device with the pins used on the Arduino sketches

### 2.4 Recording settings

The OptoPi has been tested using both the V2 and HQ Raspberry Pi camera units. Video recordings made for subsequent tracking were acquired with the Raspberry Pi HQ camera equipped with a Raspberry Pi HQ camera 6mm 3MP lens. For optogenetic experiments with IR illumination this lens can be fitted with a mounted filter that only allows the passage of infrared (IR) spectrum of the incident light (MIDOPT FIL LP830/27). The Raspberry Pi HQ camera’s IR filter can be removed according to the instructions provided by the manufacturer to make it IR-sensitive. The camera is mounted on a SG90 micro servo motor (Tower pro), rack-spur gear linear stage that converts the rotational movement of the servo motor into a linear movement (Figure 5). The distance from the camera to the arena can be adjusted within a range of approximately 50mm ([Supplementary video 4](https://osf.io/pjtmh/)).

**Figure 5.**
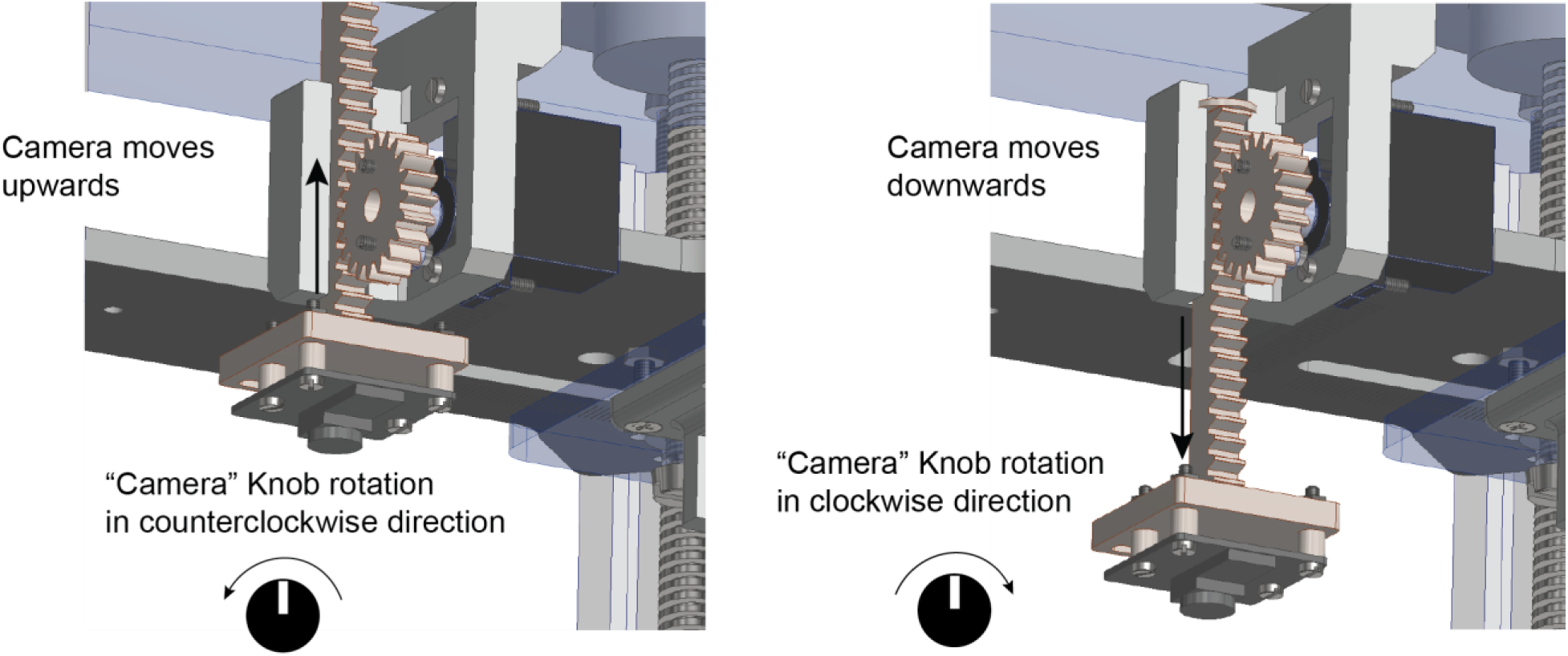
Camera vertical movement actuated from the control shield. The Camera knob actuates the servo motor of the camera translational stage moving the camera up or down and helping the user to select the right imaging distance and the right field of view.

### 2.5 Tracking options

Videos in this manuscript were acquired either using custom written python scripts on a RaspberryPi 4b running Raspbian or directly via BIO on the same RaspberryPi running Ubuntu (see Operation instructions). Videos were recorded in h264 format. Here we present a tracking software called BioImageOperation (BIO, https://joostdefolter.info/bio-research). BIO is an open source tool for real-time image processing and providing accurate tracking for downstream analysis and scientific research. It supports real-time tracking of many individuals of similar or different types, with cross-platform support including low-cost hardware.

## 3. Design files

### Design Files Summary

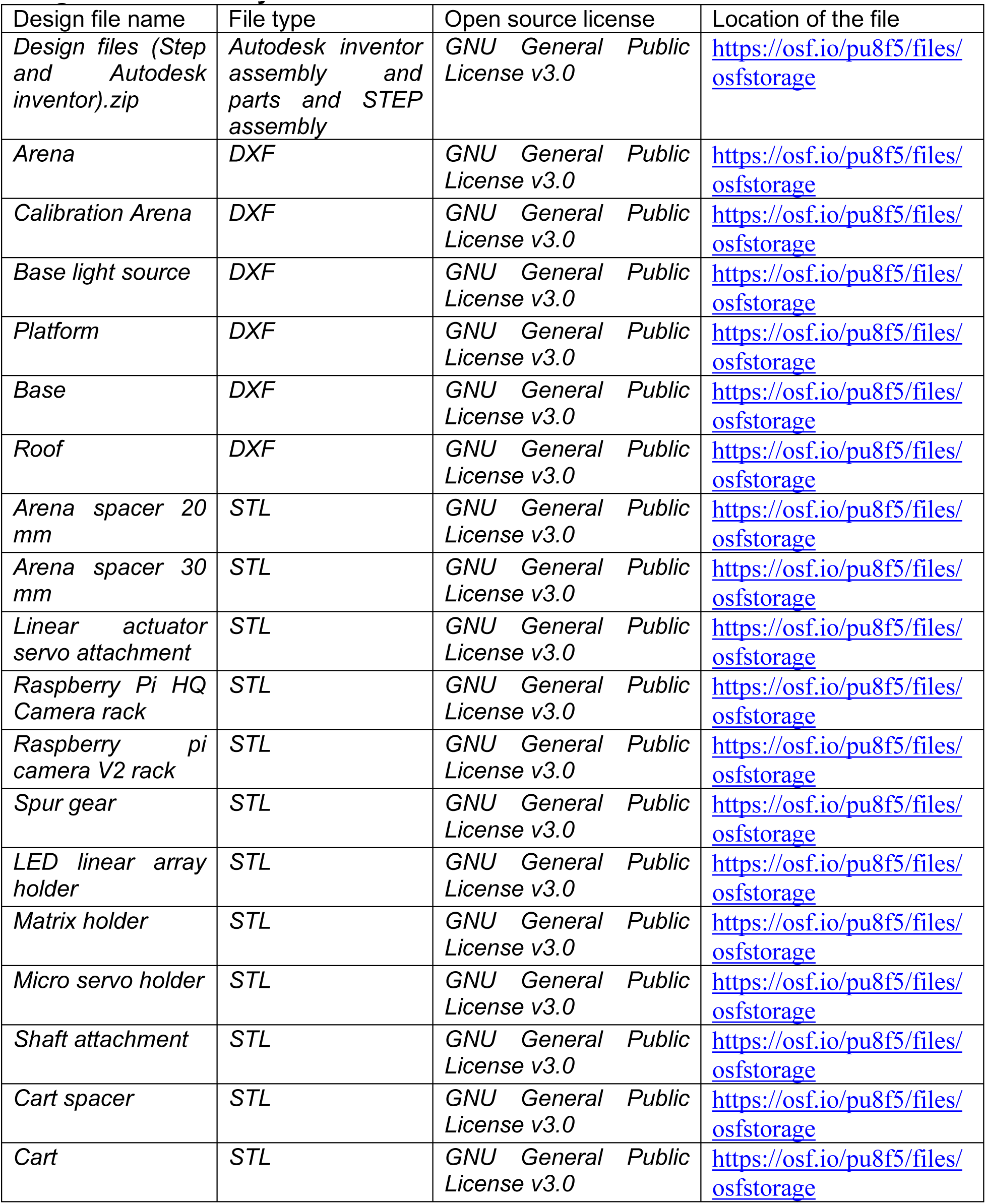
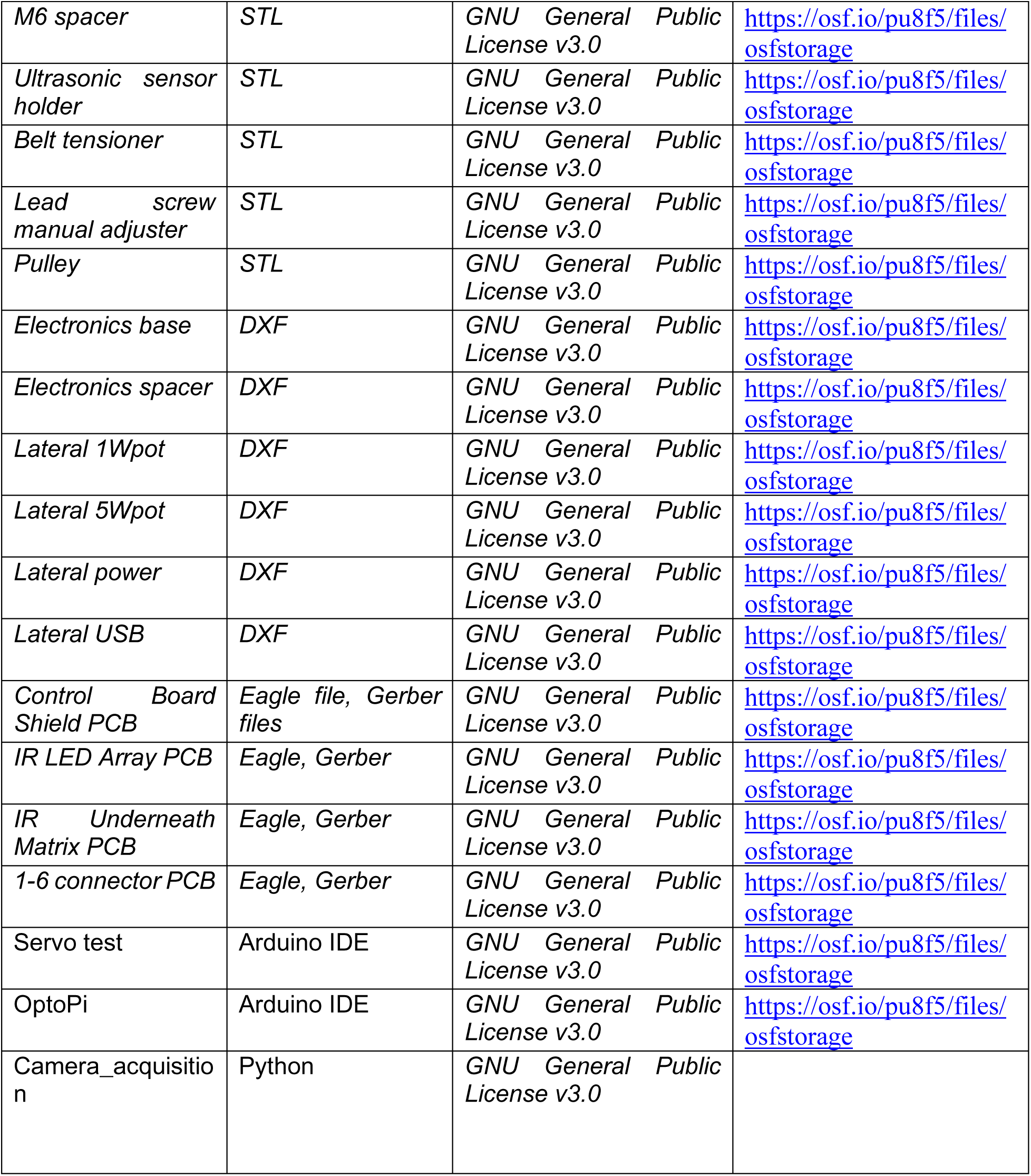

| Design file name | File type | Open source license | Location of the file |
| --- | --- | --- | --- |
| *Design files (Step and Autodesk inventor).zip* | *Autodesk inventor assembly and parts and STEP assembly* | GNU General Public License v3.0 | https://osf.io/pu8f5/files/osfstorage |
| *Arena* | *DXF* | GNU General Public License v3.0 | https://osf.io/pu8f5/files/osfstorage |
| *Calibration Arena* | *DXF* | GNU General Public License v3.0 | https://osf.io/pu8f5/files/osfstorage |
| *Base light source* | *DXF* | GNU General Public License v3.0 | https://osf.io/pu8f5/files/osfstorage |
| *Platform* | *DXF* | GNU General Public License v3.0 | https://osf.io/pu8f5/files/osfstorage |
| *Base* | *DXF* | GNU General Public License v3.0 | https://osf.io/pu8f5/files/osfstorage |
| *Roof* | *DXF* | GNU General Public License v3.0 | https://osf.io/pu8f5/files/osfstorage |
| *Arena spacer 20 mm* | *STL* | GNU General Public License v3.0 | https://osf.io/pu8f5/files/osfstorage |
| *Arena spacer 30 mm* | *STL* | GNU General Public License v3.0 | https://osf.io/pu8f5/files/osfstorage |
| *Linear actuator servo attachment* | *STL* | GNU General Public License v3.0 | https://osf.io/pu8f5/files/osfstorage |
| *Raspberry Pi HQ Camera rack* | *STL* | GNU General Public License v3.0 | https://osf.io/pu8f5/files/osfstorage |
| *Raspberry pi camera V2 rack* | *STL* | GNU General Public License v3.0 | https://osf.io/pu8f5/files/osfstorage |
| *Spur gear* | *STL* | GNU General Public License v3.0 | https://osf.io/pu8f5/files/osfstorage |
| *LED linear array holder* | *STL* | GNU General Public License v3.0 | https://osf.io/pu8f5/files/osfstorage |
| *Matrix holder* | *STL* | GNU General Public License v3.0 | https://osf.io/pu8f5/files/osfstorage |
| *Micro servo holder* | *STL* | GNU General Public License v3.0 | https://osf.io/pu8f5/files/osfstorage |
| *Shaft attachment* | *STL* | GNU General Public License v3.0 | https://osf.io/pu8f5/files/osfstorage |
| *Cart spacer* | *STL* | GNU General Public License v3.0 | https://osf.io/pu8f5/files/osfstorage |
| *Cart* | *STL* | GNU General Public License v3.0 | https://osf.io/pu8f5/files/osfstorage |
| *M6 spacer* | *STL* | *GNU General Public License v3.0* | https://osf.io/pu8f5/files/osfstorage |
| *Ultrasonic sensor holder* | *STL* | *GNU General Public License v3.0* | https://osf.io/pu8f5/files/osfstorage |
| *Belt tensioner* | *STL* | *GNU General Public License v3.0* | https://osf.io/pu8f5/files/osfstorage |
| *Lead screw manual adjuster* | *STL* | *GNU General Public License v3.0* | https://osf.io/pu8f5/files/osfstorage |
| *Pulley* | *STL* | *GNU General Public License v3.0* | https://osf.io/pu8f5/files/osfstorage |
| *Electronics base* | *DXF* | *GNU General Public License v3.0* | https://osf.io/pu8f5/files/osfstorage |
| *Electronics spacer* | *DXF* | *GNU General Public License v3.0* | https://osf.io/pu8f5/files/osfstorage |
| *Lateral 1Wpot* | *DXF* | *GNU General Public License v3.0* | https://osf.io/pu8f5/files/osfstorage |
| *Lateral 5Wpot* | *DXF* | *GNU General Public License v3.0* | https://osf.io/pu8f5/files/osfstorage |
| *Lateral power* | *DXF* | *GNU General Public License v3.0* | https://osf.io/pu8f5/files/osfstorage |
| *Lateral USB* | *DXF* | *GNU General Public License v3.0* | https://osf.io/pu8f5/files/osfstorage |
| *Control Board Shield PCB* | *Eagle file, Gerber files* | *GNU General Public License v3.0* | https://osf.io/pu8f5/files/osfstorage |
| *IR LED Array PCB* | *Eagle, Gerber* | *GNU General Public License v3.0* | https://osf.io/pu8f5/files/osfstorage |
| *IR Underneath Matrix PCB* | *Eagle, Gerber* | *GNU General Public License v3.0* | https://osf.io/pu8f5/files/osfstorage |
| *1-6 connector PCB* | *Eagle, Gerber* | *GNU General Public License v3.0* | https://osf.io/pu8f5/files/osfstorage |
| *Servo test* | *Arduino IDE* | *GNU General Public License v3.0* | https://osf.io/pu8f5/files/osfstorage |
| *OptoPi* | *Arduino IDE* | *GNU General Public License v3.0* | https://osf.io/pu8f5/files/osfstorage |
| *Camera_acquisition* | *Python* | *GNU General Public License v3.0* |  |

## 4. Bill of Materials

### Bill of Materials

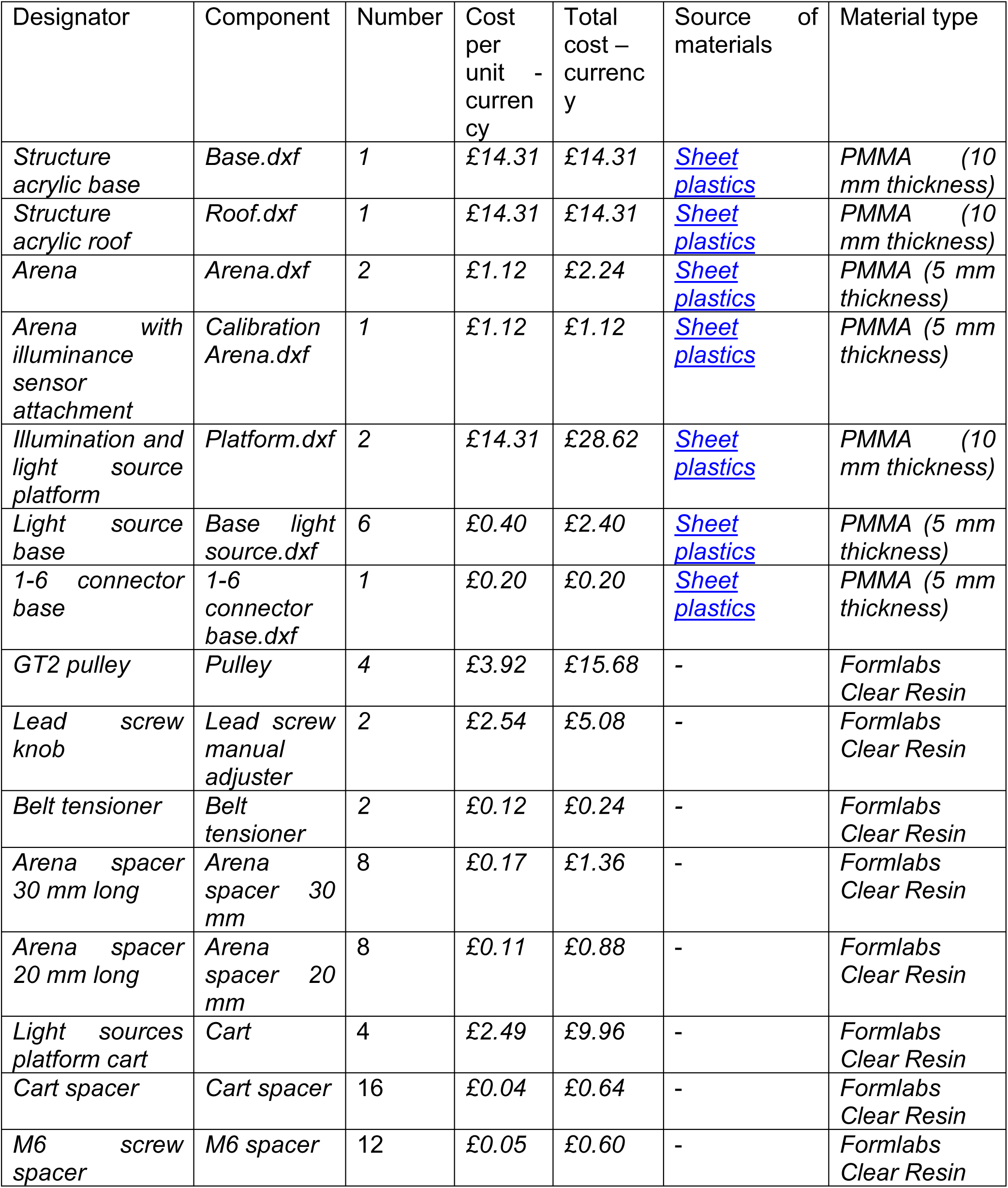
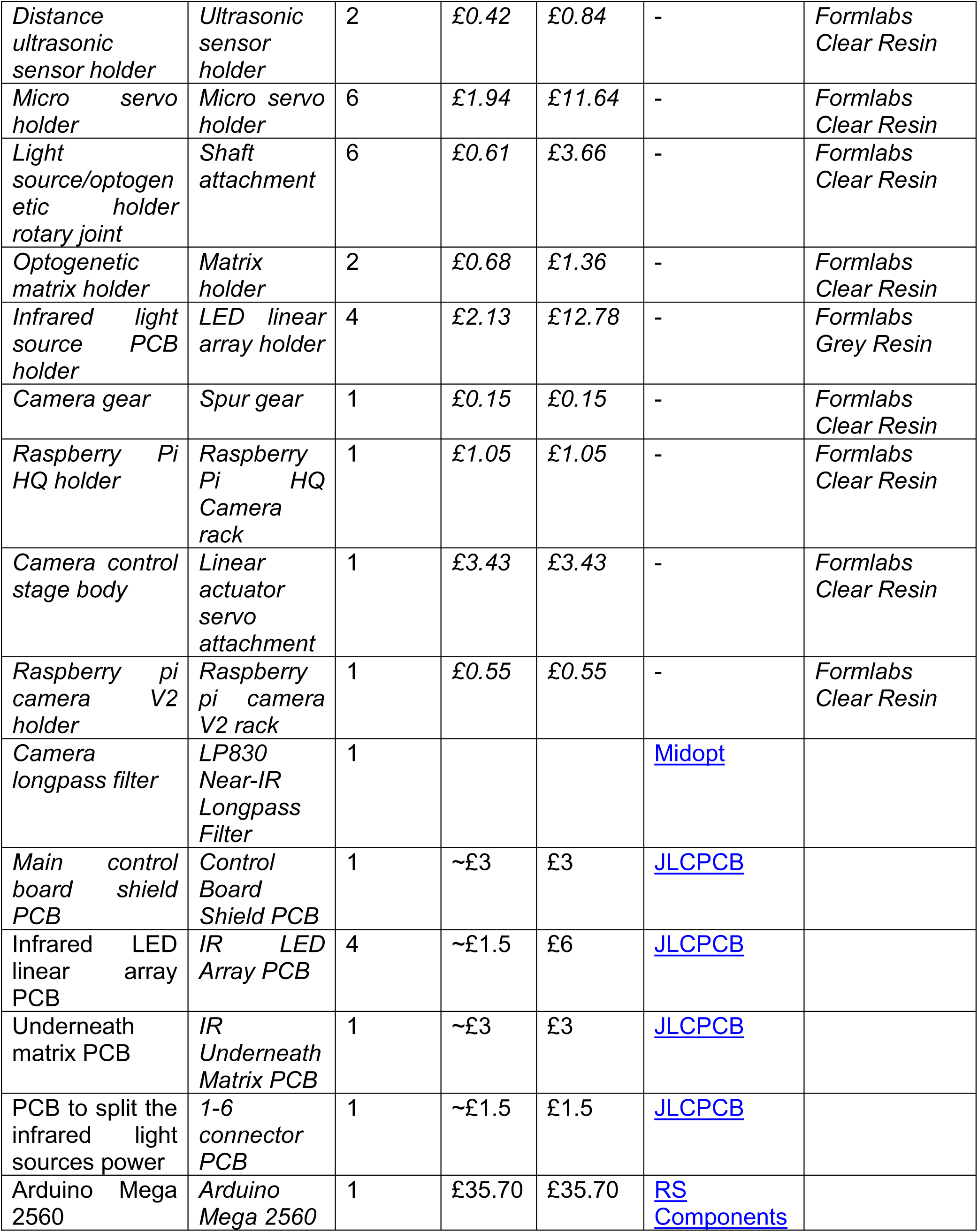
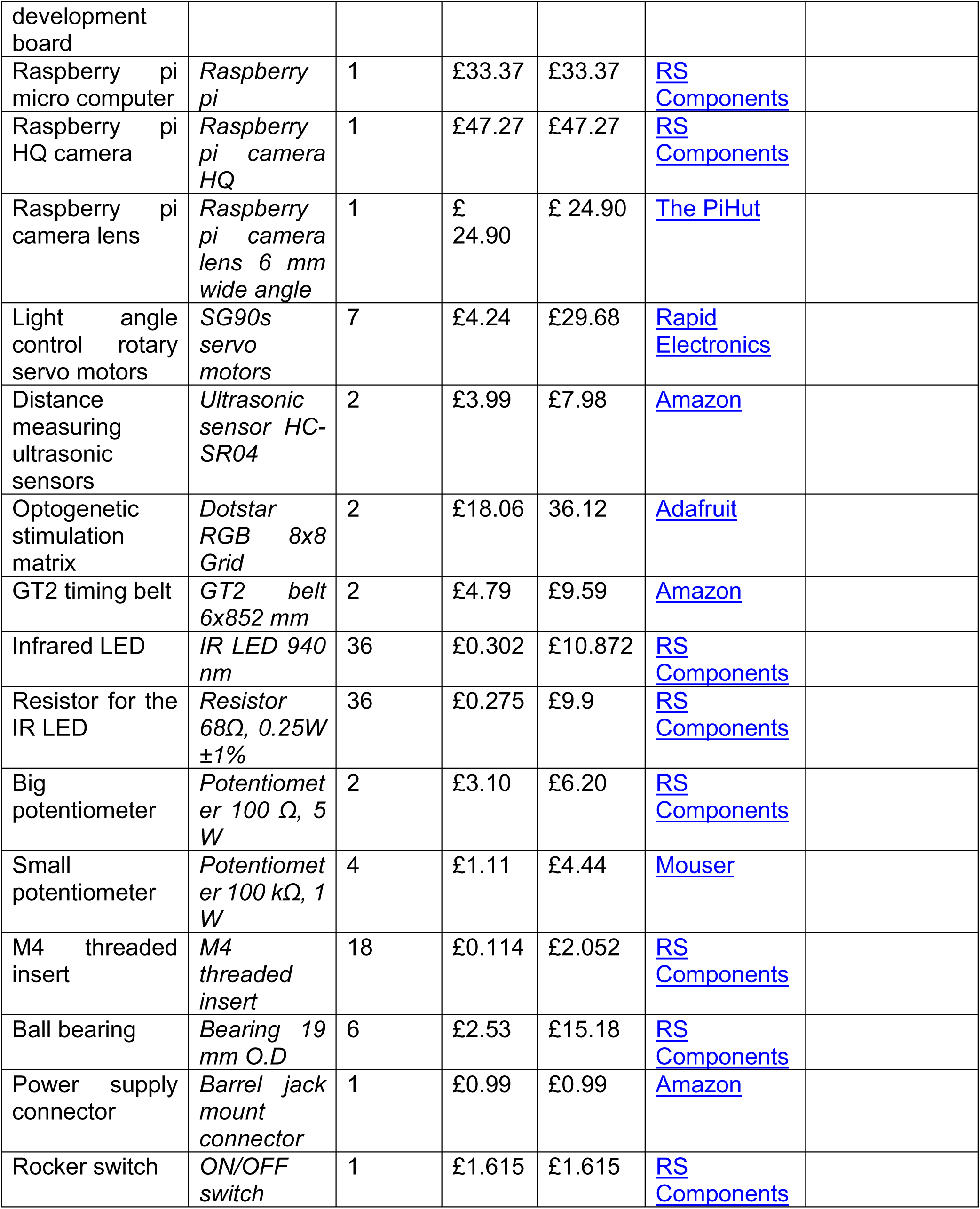
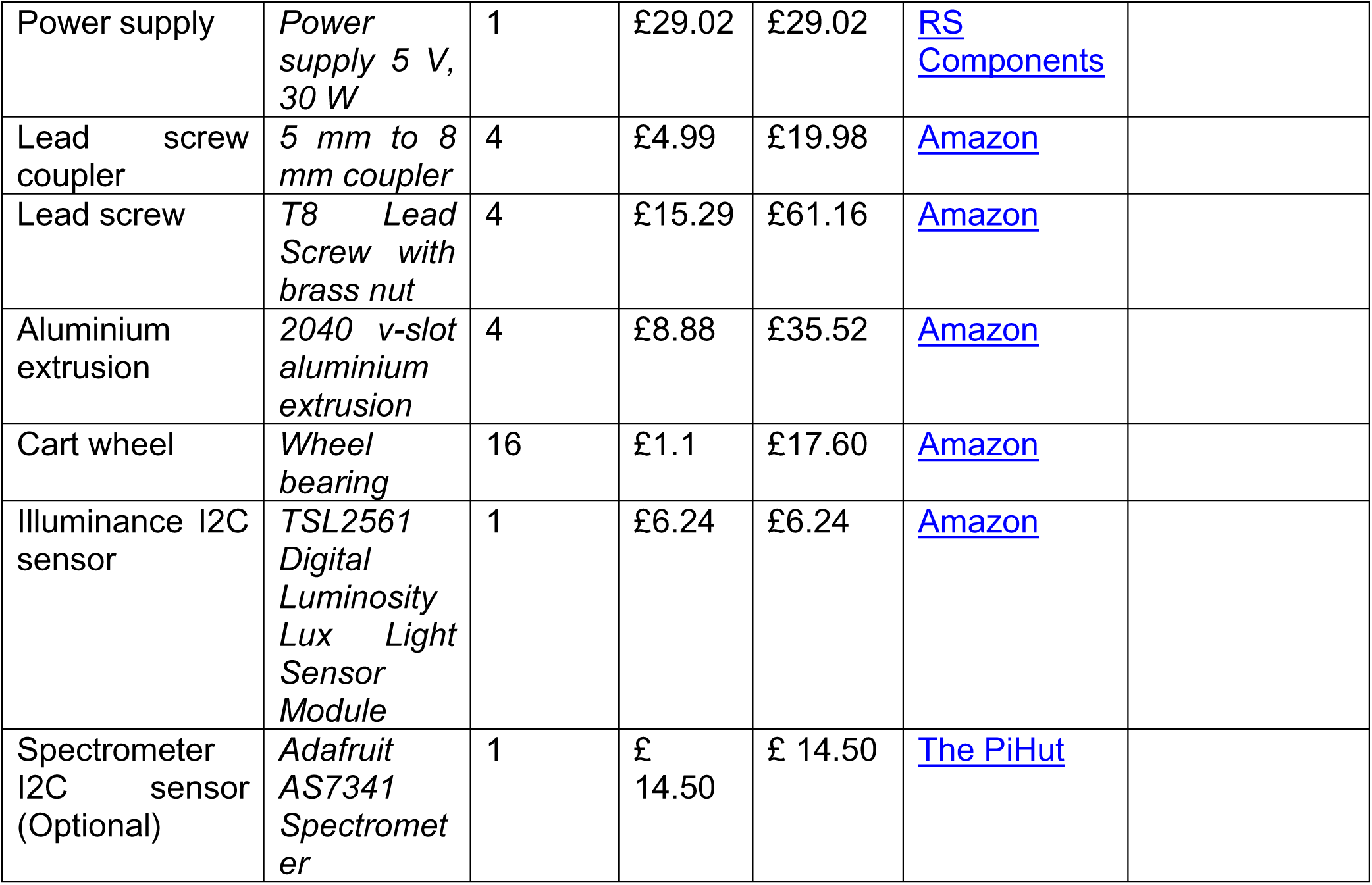

## 5. Build Instructions

This section has the following subsections:

5.1. Generate all mechanical pieces
5.2. Assembly of the illumination infrared LED arrays
5.3. Assembly of the RGB Matrix holder
5.4. Camera linear stage assembly
5.5. Arena assembly
5.6. Cart assembly
5.7. Platform assembly
5.8. Ultrasonic sensors assembly
5.9. Device final mechanical assembly
5.10. Soldering
5.11. Electrical connections

### 5.1. Generate all mechanical pieces

1- Print all required 3D printed pieces (Figure 6). In our case we used a Formlabs Form 3 3D printer with 100 µm resolution and Clear and Gray resins.

**Figure 6.**
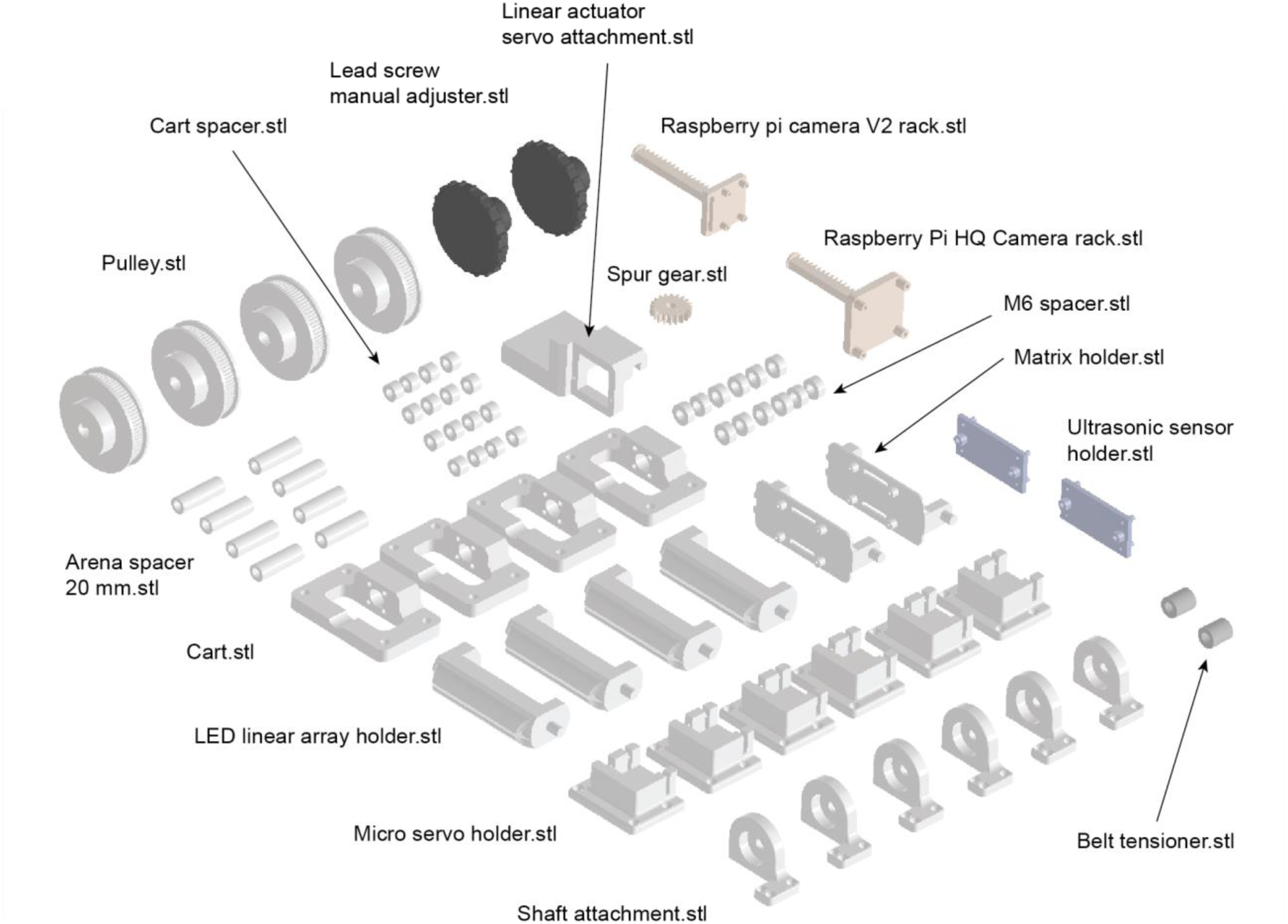
OptoPi components to 3D print.

2- Laser cut all of the required pieces (Figure 7 and 8). In our case we used a Universal laser systems VLS3.50 laser cutter and PMMA sheets (5 and 10 mm thickness).

**Figure 7.**
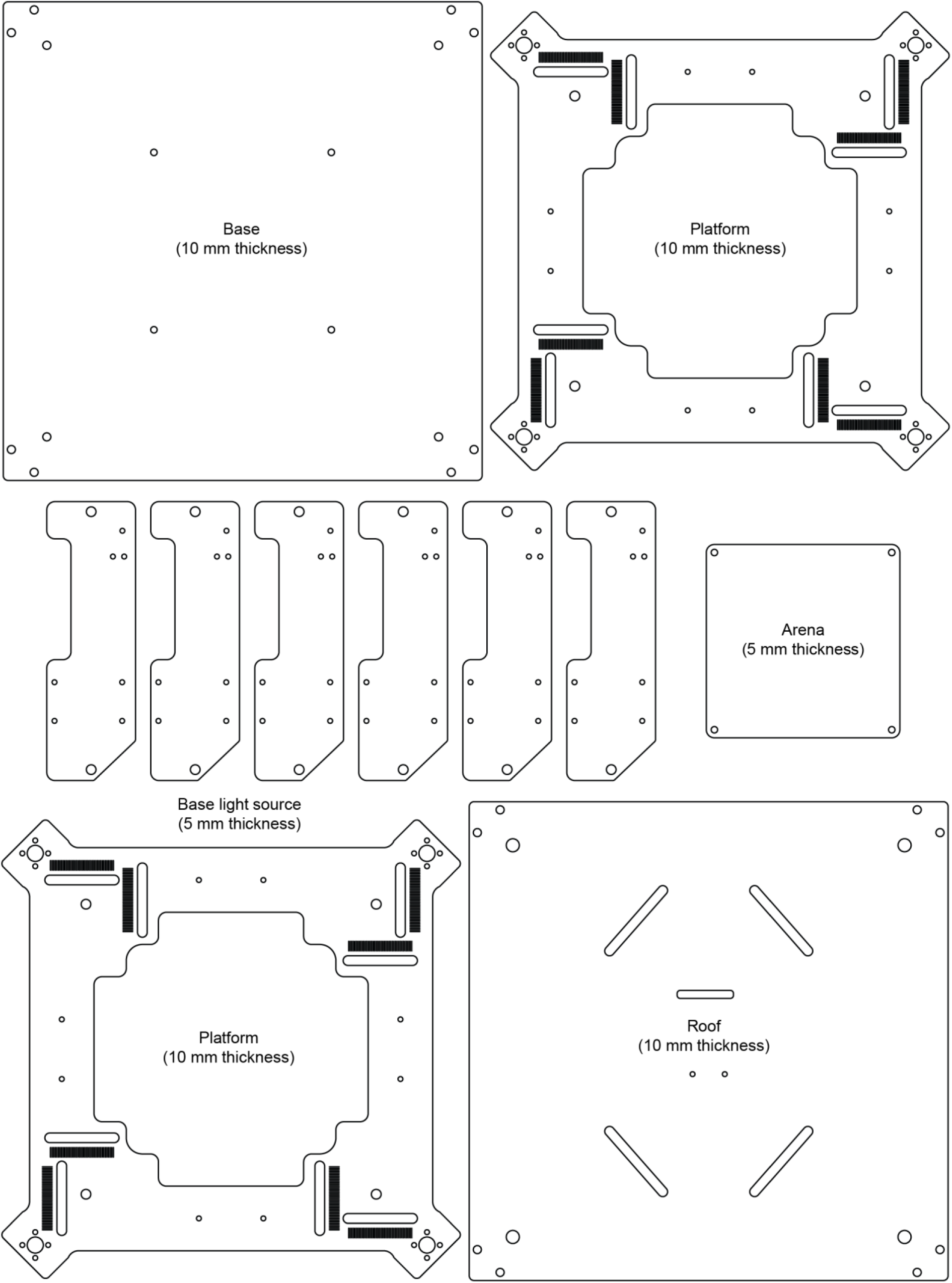
OptoPi components to laser cut.

**Figure 8.**
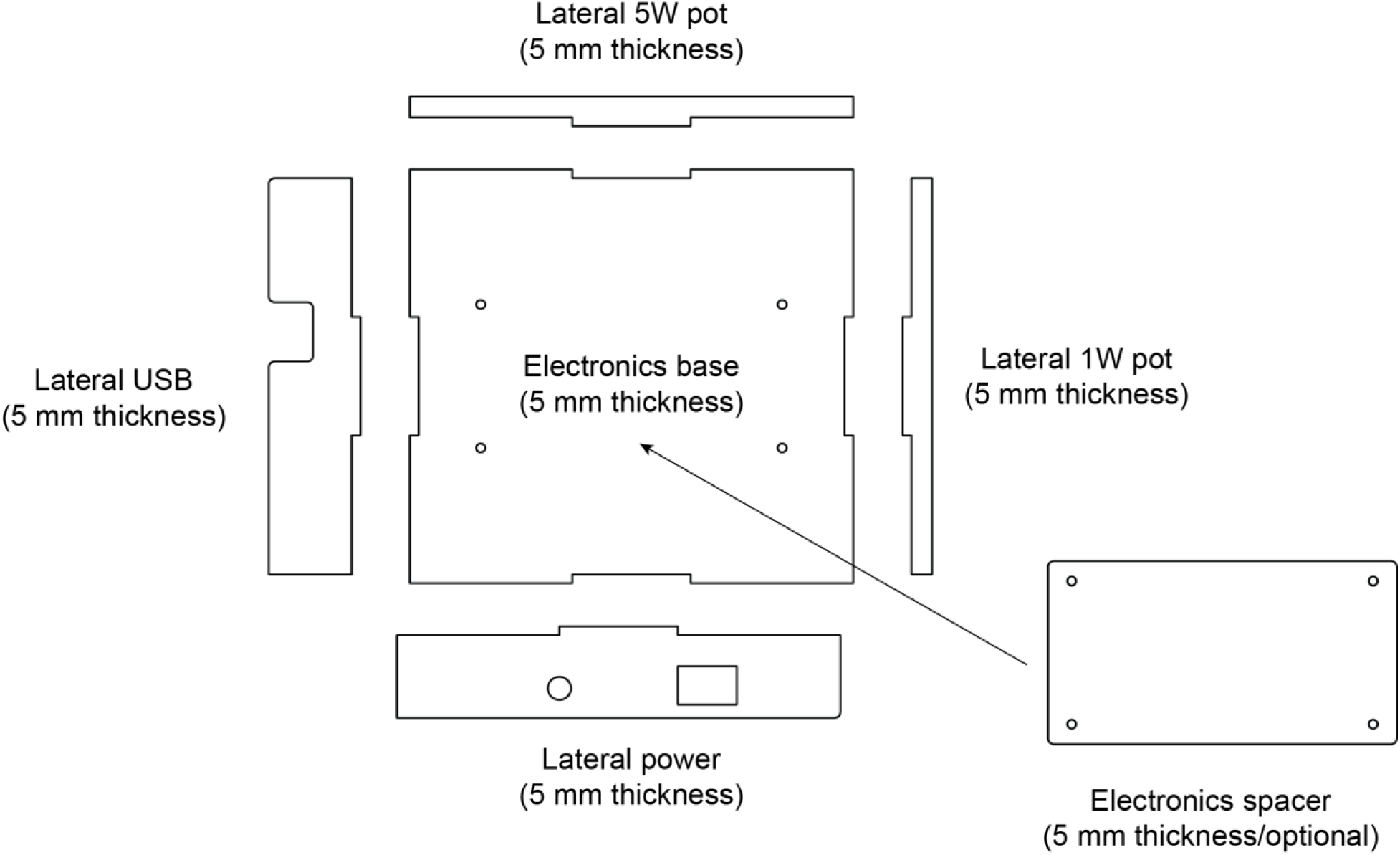
Electronics control box components to laser cut.

### 5.2. Assembly of the illumination infrared LED arrays

1. Insert the ball bearing on the Shaft attachment. The shaft of the LED linear array holder fits on the bearing just by push fit (Figure 9c).
2. Attach the micro servo on the micro servo holder with two M2 screws (Figure 9d).
3. Insert the two M2 screws on the servo arm (Figure 5c).
4. Connect the servo motor to the Arduino 5V, GND and Pin 9 (Figure 9e).
5. Upload the program *Servo_test* to the Arduino board.
6. Connect the Arduino with the USB cable.
7. Unscrew the Servo arm.
8. Place the servo arm vertical (Figure 9f) The vertical position can vary slightly from one servo motor to the other. This can be adjusted afterwards adding or subtracting one degree to the particular servo motor that is not aligned with the others.
9. Do the same with all the servo motors of the setup that are moving RGB LED matrices or IR LED arrays.
10. All the steps have to be done for the seven servo motors attached to their holders.
11. Then insert two M2 screws on the servo white arm (Figure 9g).
12. Screw this two M2 screws with the LED holder (Figure 9h).
13. Solder the IR LED 940 nm and the 68Ω resistors on the LED Array PCB (Figure 9i).
14. Insert the LED array on their holder and insert 4 screws to hold the PCB (Figure 9j).
15. Attach the previous assembly with the light source base laser cut part (Figure 9k).
16. Repeat the process for each of the four LED linear array.

**Figure 9.**
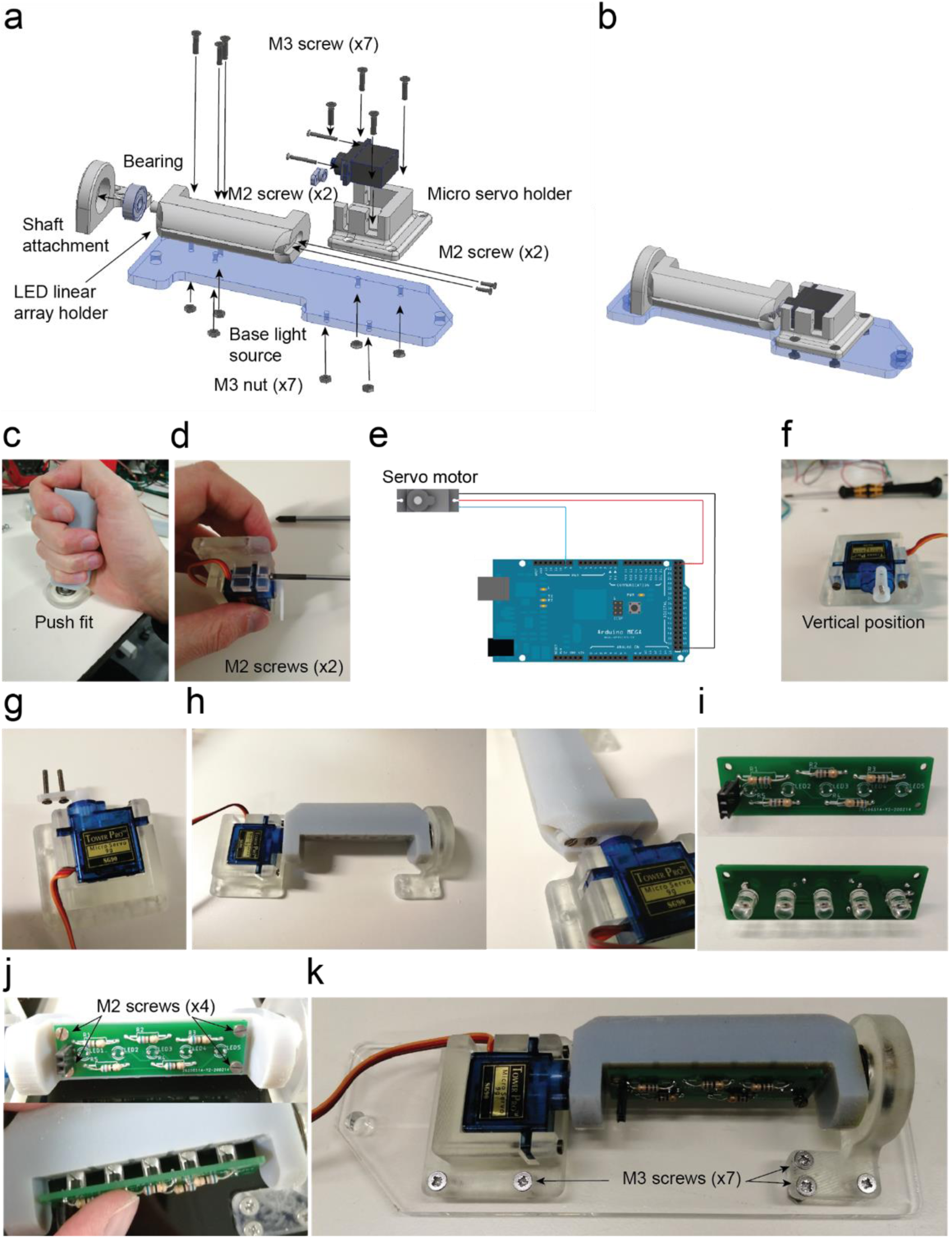
Infrared LED array assembly. **a.** Exploded view of the illumination infrared LED arrays. **b.** Assembled view of the illumination infrared LED arrays ([Supplementary video 5](https://osf.io/ja9rc/)). **c.** LED linear array holder inserted into the bearing. **d.** Servo motor attached to holder with two M2 screws. **e.** SG90s servo motor connection. **f.** Position of the servo’s arm required for the calibration. **g.** Two M2 screws are placed on the servo lever after drilling two holes to increase their diameter. **h.** Assembly completed and detail of the two M2 screws threaded into the LED linear array holder part. **i.** Infrared LED soldered on the LED array PCB. 68Ω resistors are soldered onto the other side of the PCB which is attached with four M2 screws threaded on the plastic. **j.** The PCB is attached with 4 M2 screws on the 3D printed part. The LEDs are inserted into five holes by pushing them through. **k.** The assembly is completed by attaching it on the base light source part with seven M3 screws and their respective nuts.

### 5.3. Assembly of the RGB Matrix holder

1. Insert the ball bearing on the Shaft attachment (Figure 10a-c).
2. Insert the shaft of the RGB LED Matrix holder on the bearing hole leaving a 0.5 mm distance to avoid friction (Figure 10c-d).
3. Attach the micro servo on the micro servo holder with two M2 screws as done with the Infrared holders before.
4. Insert the two M2 screws on the servo arm.
5. Screw thes two screws with the LED holder (Figure 10e).
6. Solder the 220 µF capacitor and the Vcc, GND, Data and Clock wires on the matrix pads (Figure 10f).
7. Attach the Matrices on the holder with four M2 screws (Figure 10f).
8. Attach the system with the Base light source (Figure 10g).

**Figure 10.**
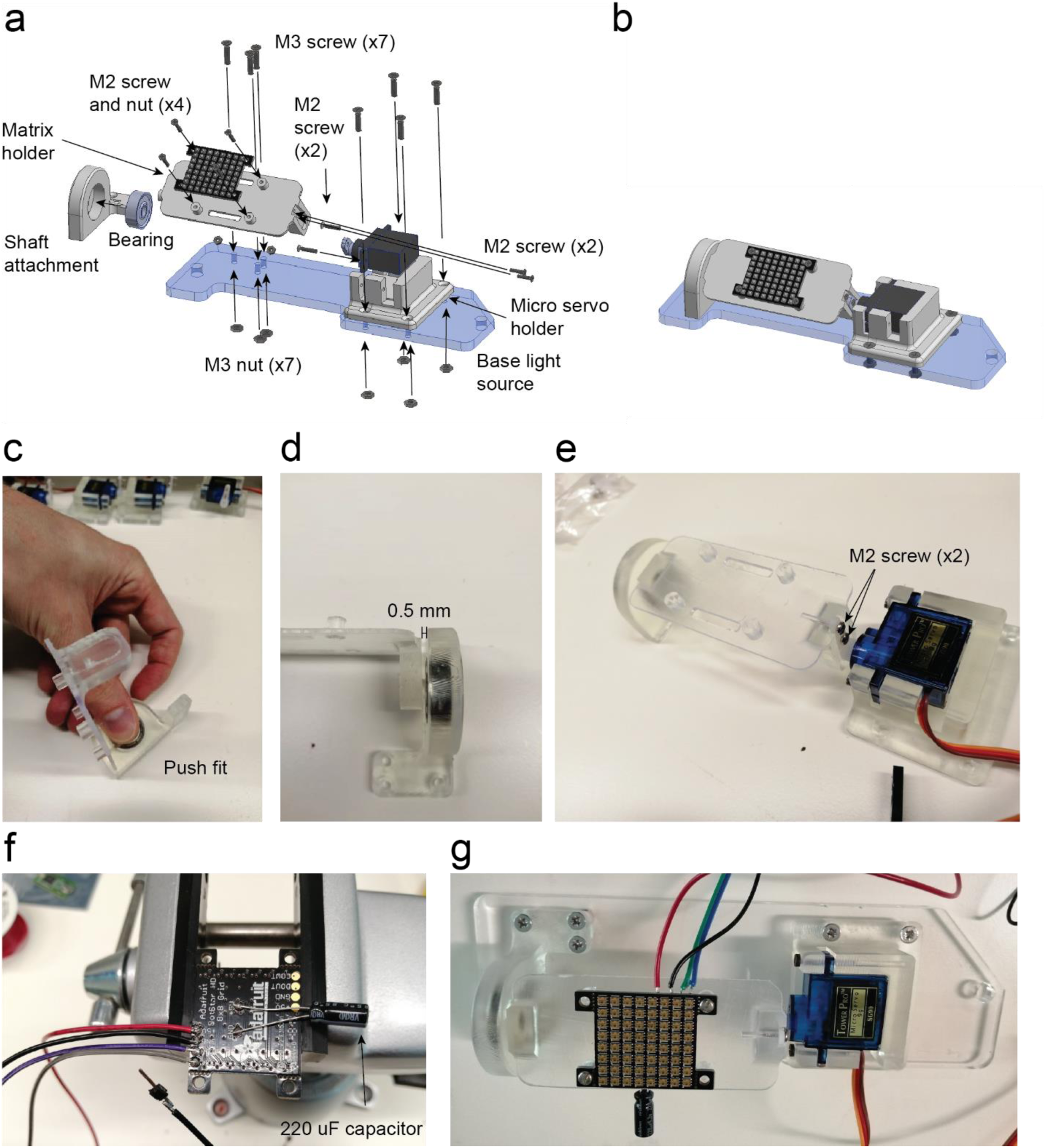
RGB matrix holder assembly. **a.** Exploded view of the RGB matrix holder. **b.** Assembled view of the RGB matrix holder ([Supplementary video 6](https://osf.io/ha2dn/)). **c.** The shaft of the matrix holder fits on the bearing by push fit. **d.** Assembled RGB matrix holder with the detail in the space between the bearing and the 3D printed part to avoid friction. **e.** Assembled RGB matrix with two M2 screws. **f.** Detail in the wires soldered on the RGB grid pads as well as the 220 µF capacitor that comes with the grid. **g.** The assembly is completed by attaching it onto the base light source part with seven M3 screws and their respective nuts.

### 5.4. Camera linear stage assembly

1. Cut the servo motor white adaptor which has two symmetric sides leaving just the same diameter as the internal diameter of the spur gear (Figure 11c-e).
2. Insert the two M3 threaded inserts with the linear actuator servo attachment just applying pressure manually (Figure 11f).
3. Attach the servo motor with the linear actuator servo attachment using two M2 screws and nuts (Figure 11g-h).
4. Place the rack on the guide and the spur gear with their teeth coincident and use the servo motor shaft screw to attach the spur gear with the servo motor. Make sure the servo motor has the whole rotational range in a useful linear range of motion of the rack (Figure 11i-j).

**Figure 11.**
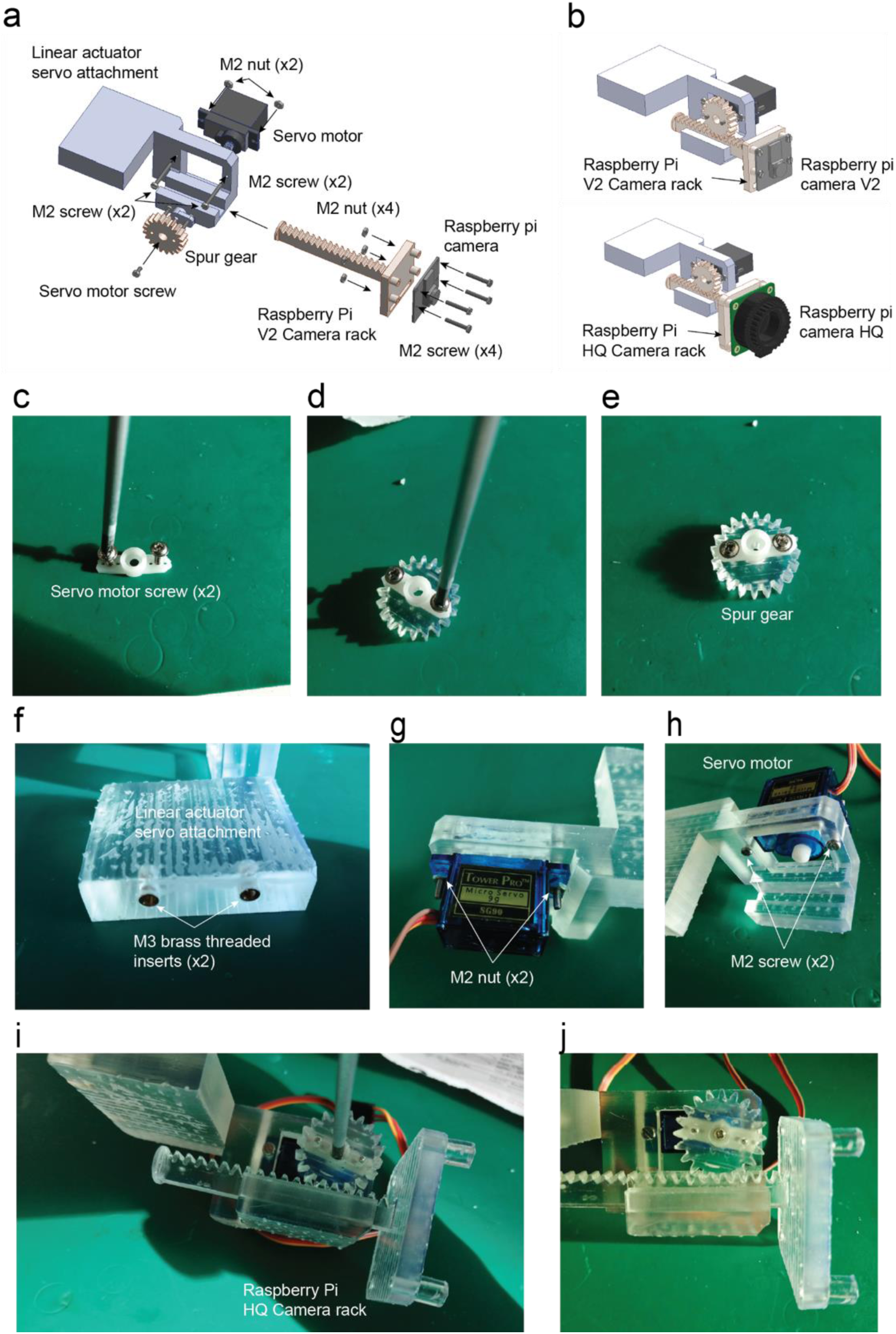
Camera linear stage assembly. **a.** Exploded view. **b.** Assembled view ([Supplementary video 7](https://osf.io/he796)). **c.** The servo motor arm is shortened by cutting both sides at the level of the third hole. Then the servo motor screws are screwed in the second hole on each side of the servo motor. **d.** The arm is placed over the spur gear with the holes coincident. **e.** The servo arm and the spur gear are attached with the screws. **f.** Two M3 threaded inserts are inserted by applying pressure before curing the resin. **g.** The servo motor is attached using two M2 screws with M2 nuts. **h.** Detail of the M2 screws on the front view of the assembly. **i.** Spur gear assembly, the rack is placed in the channel and then the spur gear is attached making the teeth coincident and keeping the maximum travel range. **j.** The servo motor screw holds the spur gear as a final step.

### 5.5. Arena assembly

1. Use M4 grub screws for each level of the arena you want and regular M4 screws for the top (Figure 15a).
2. Laser cut the Arena and 3D print the spacers. The arena can be laser cut with different materials with different optical properties to get diffusivity or opacity of the matrix placed underneath (Figure 12c).
3. Insert in every 3D printed arena spacer M4 threaded inserts on each side (Figure 15d).

**Figure 12.**
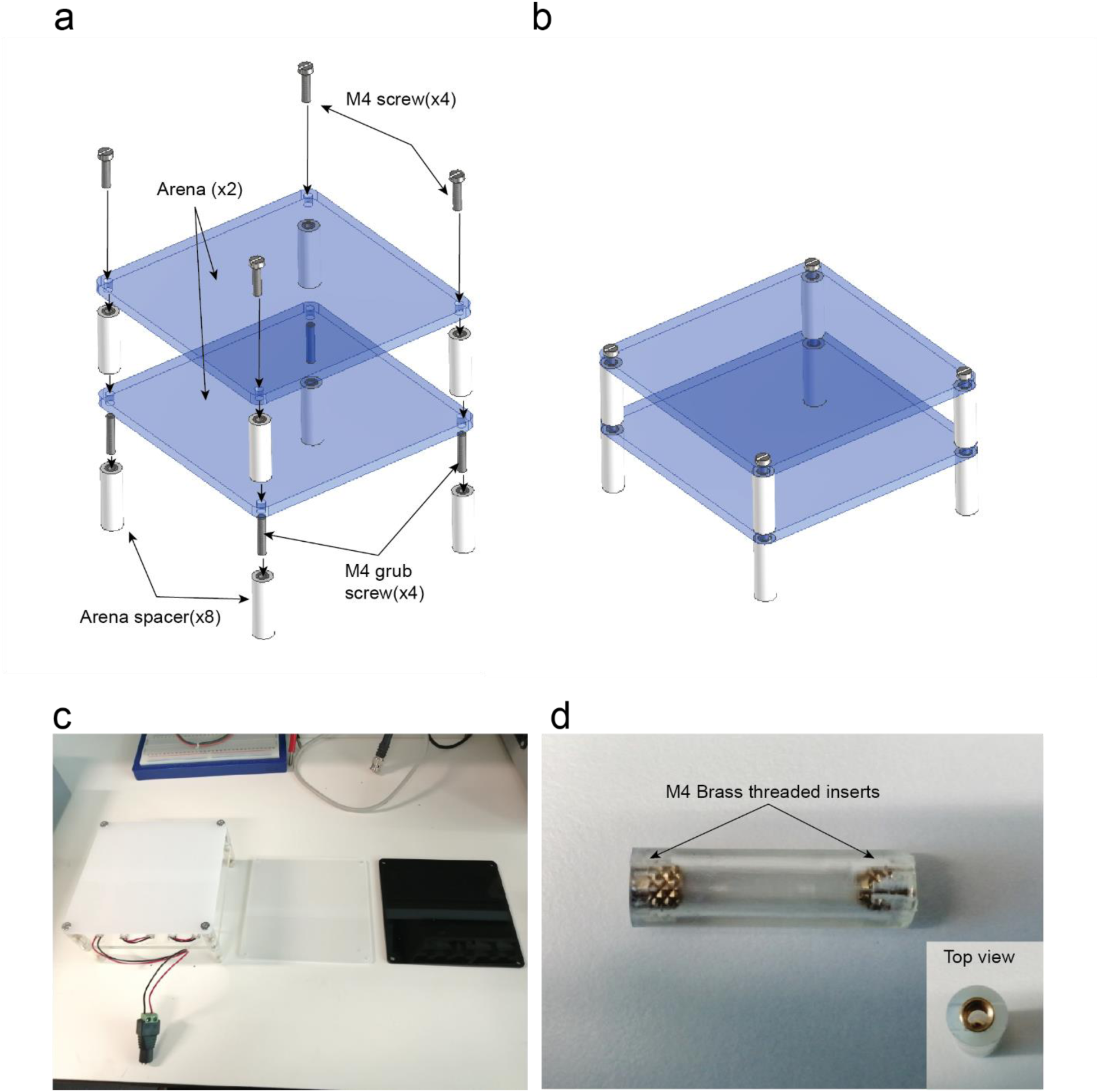
Arena assembly. **a.** Exploded view. **b.** Assembled view ([Supplementary video 8](https://osf.io/ua4k9)). **c.** Different laser cut arena sheets using materials with different opacities. **d.** Lateral view of the spacer with threaded inserts and top view of the spacer

### 5.6. Cart assembly

1. Attach the wheels with the cart placing a cart spacer in between and a M5 screw in between (Fig 13a-b).
2. Insert the lead screw nut on the central hole of the cart (Fig 13a-b).
3. Attach two carts with the platform using four long M3 screws. The screws are attached to the lead screw nut (Fig 13c-d).

**Figure 13.**
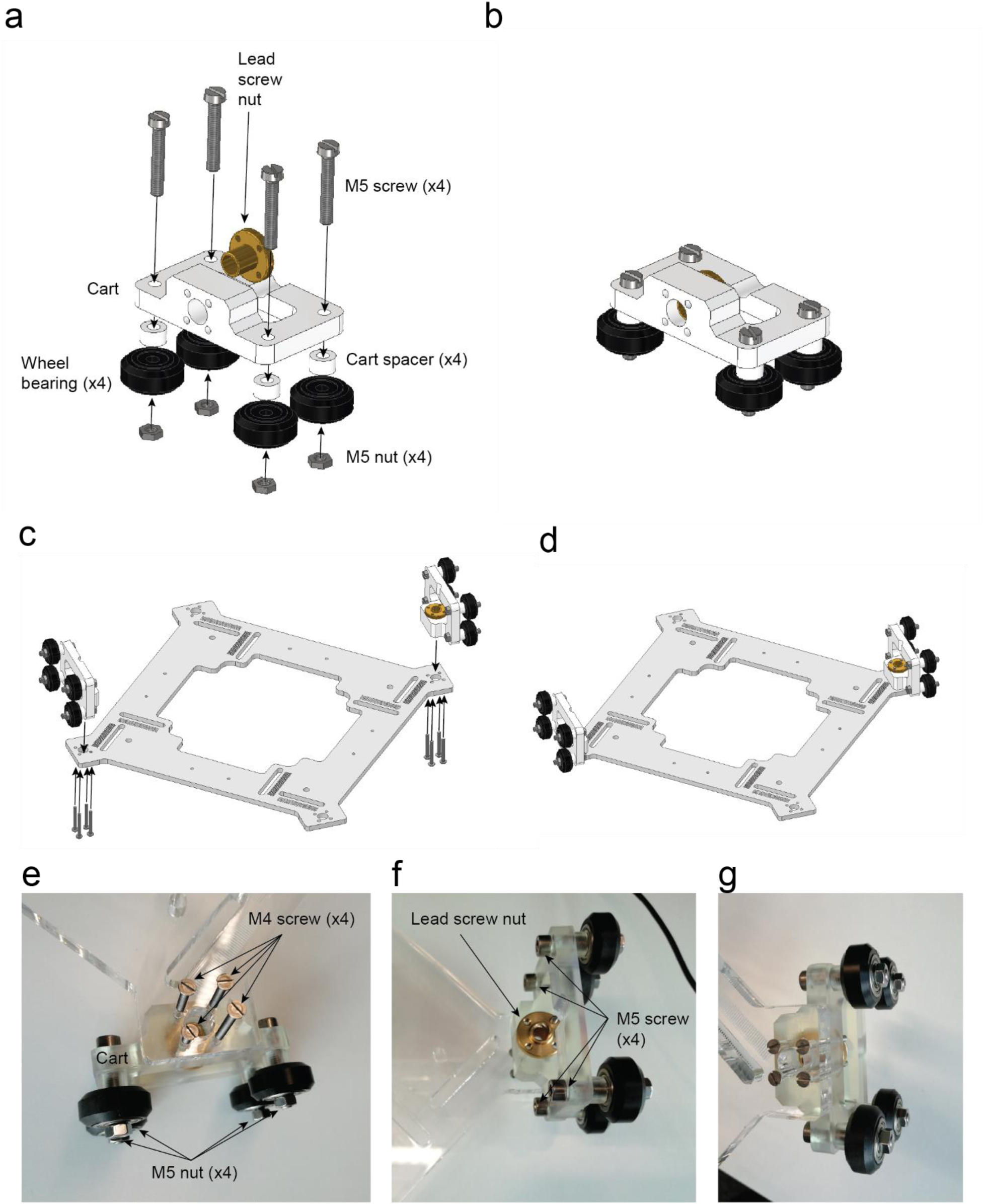
Cart assembly. **a.** Exploded view. **b.** Assembled view ([Supplementary video 9](https://osf.io/g8sp4)). Platform assembly. **c.** Exploded view of the platform assembly. **d.** Platform assembled with both carts ([Supplementary video 10](https://osf.io/jc95g)). **e.** Lead screw nut. **f.** Long M3 screws (50 mm) are needed to hold the nut in place. **g.** Detail of the four M3 screws.

### 5.7. Platform assembly

1. Using M6 screws, M6 spacers and M6 nuts the four IR light sources previously assembled can be now attached to the platform (Fig 14a-b).
2. The same applies for the two RGB matrices (Fig 14c-d).

**Figure 14.**
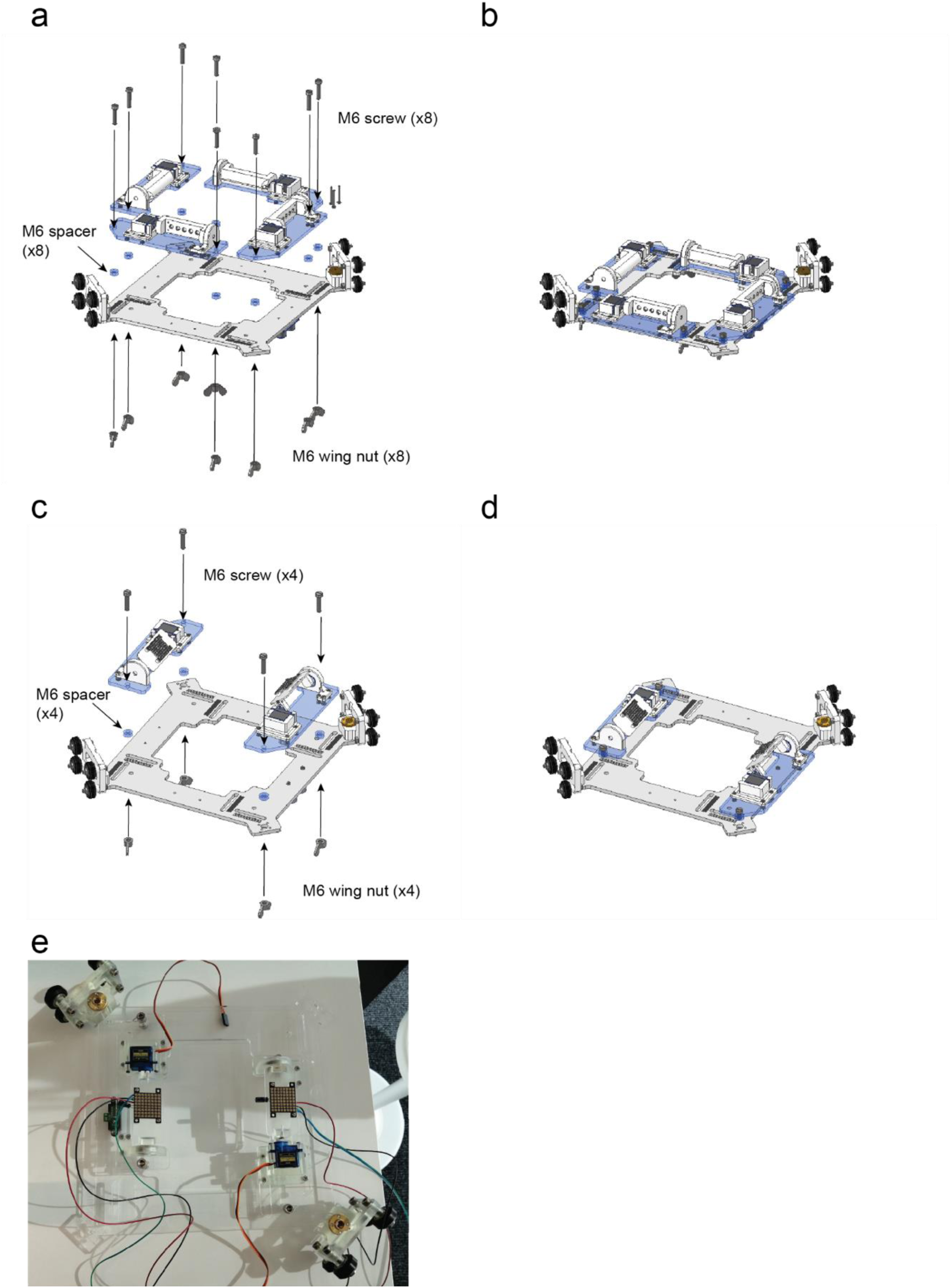
Platform with IR light sources assembly. **a.** Exploded view. **b.** Assembled view ([Supplementary video 11](https://osf.io/gbupx)). Platform with RGB light sources assembly. **c.** Exploded view. **d.** Assembled view ([Supplementary video 12](https://osf.io/5z4sh)). **e.** RGB light sources assembled with the platform.

### 5.8. Ultrasonic sensors assembly

1. Attach the Ultrasonic sensor holder with the platform using two M3 screws and two M3 nuts (Figure 15a).
2. Then attach the Ultrasonic sensor with the holder using four M1.6 screws (Figure 15a-d).
3. The M1.6 screws are inserted directly in the plastic (Figure 15c-d).

**Figure 15.**
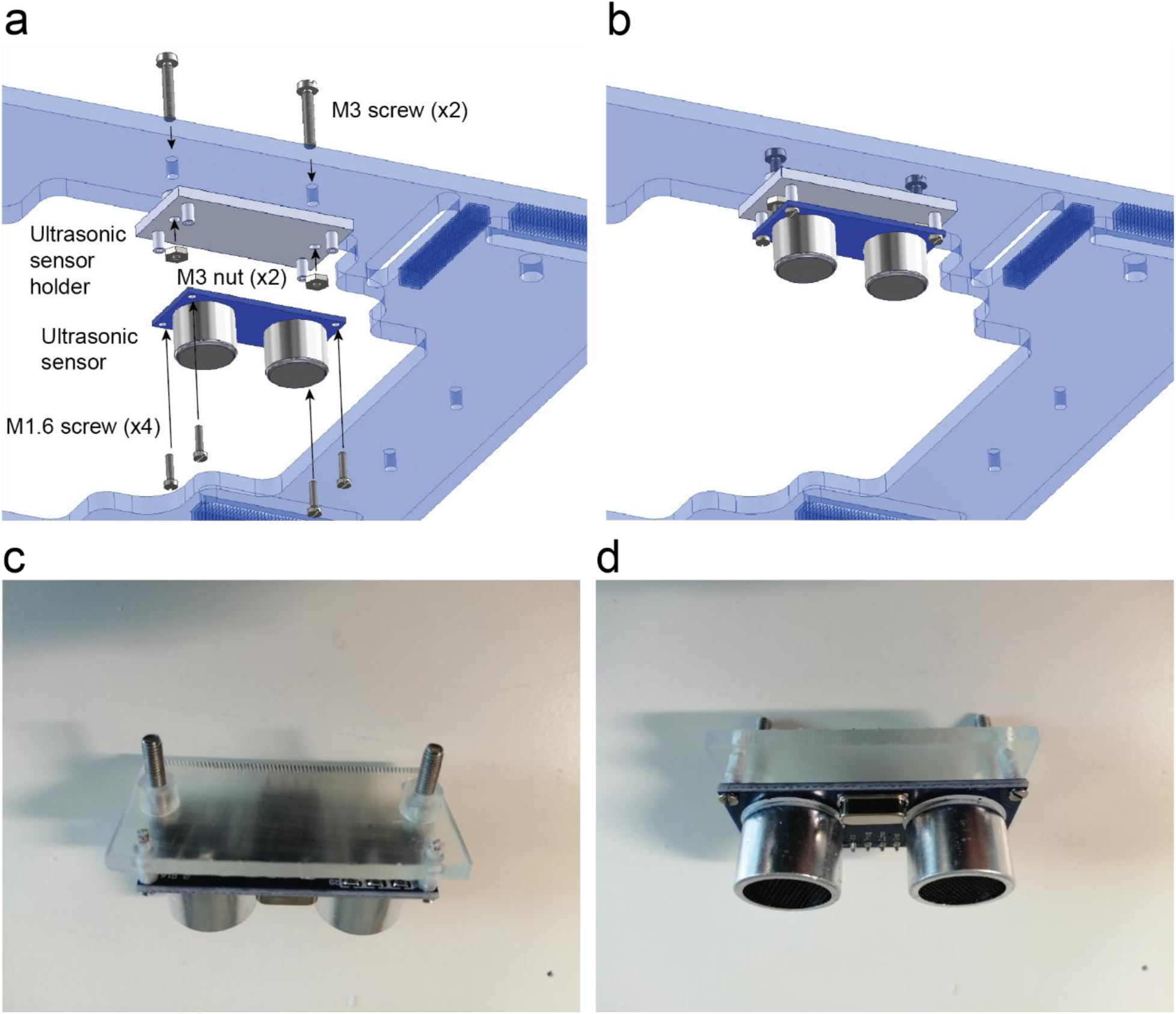
Ultrasonic sensor assembly. **a.** Exploded view. **b.** Ultrasonic sensor assembled ([Supplementary video 13](https://osf.io/5z4sh/)). **c.** Back view. **d.** Front view.

### 5.9. Device final mechanical assembly

1. Use two M3 screws to attach the camera stage with the roof (Figure 16a).
2. Attach the roof with the structure using eight more M5 screws (Figure 16a).
3. Put the four pulleys on their respective lead screws keeping one diagonal higher than the other one, at least the thickness of the belt (6mm). Use an M4 grub screw for each pulley (Figure 16a).
4. Attach the arena to the base using four M4 screws (Figure 16b).
5. The lead screws can be now inserted to the four couplers and tightened using the radial M4 grub screws of the coupler (Figure 16b).
6. The two platforms can be inserted by rotating manually the lead screws and making sure that the platform does not tilt (Figure 16b).
7. The couplers can also be attached to the base using M5 screws (Figure 16f).
8. The four aluminium profile holes have to be threaded with an M5 tap tool (RS Components 917-2599) (Figure 16d).
9. Then the aluminium extrusions can be attached to the base with M5 screws (Figure 16e).
10. Place the two 3D printed manual adjusters on the front lead screws and tighten their M4 grub screws (Figure 16a – 16g).
11. Place the GT2 timing belts and add the 3D printed belt tensioners using the diagonal guides designed for this purpose. Use M6 screws and nuts to keep them vertical and confined between two M6 nuts (Figure 16g).

**Figure 16.**
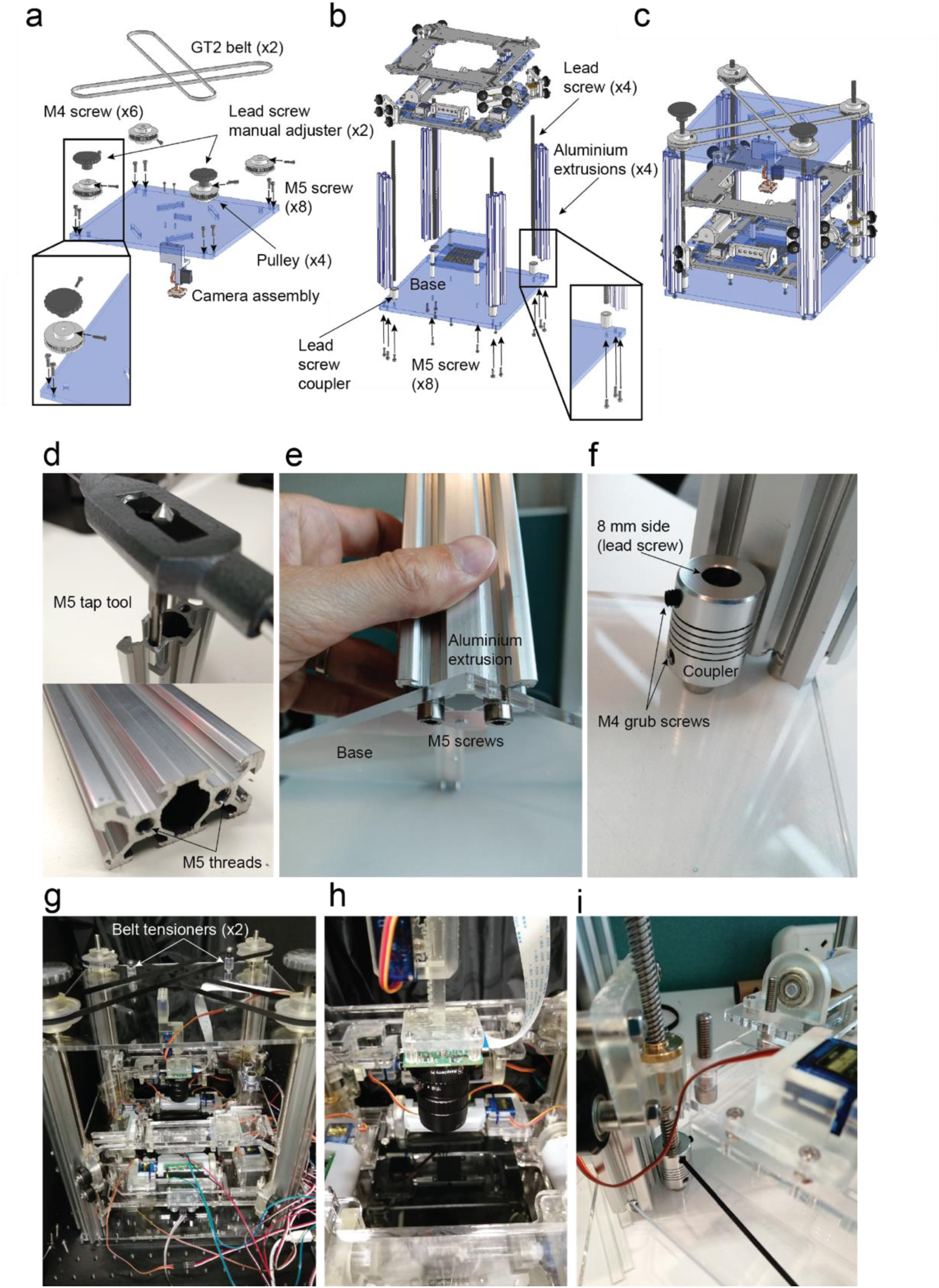
OptoPi final assembly. **a.** Belt system assembly. **b.** Assembly of aluminium extrusions 2040, lead screws, arena and both illumination platforms. **c.** Device assembled ([Supplementary video 14](https://osf.io/bacyz/), 15, 16). Aluminium extrusions and lead screw coupler details. **d.** Tap tool M5 size and M5 treads after the process. **e.** The aluminium 2040 v-slot extrusions are attached with M5 screws to the base after threading the holes they have on both sides. **f.** The lead screw coupler is held by a M5 screw placed through the base. Assembly of the main structure. **g.** Detail of the timing belt system and how the belt tensioner are confined from the vertical displacement using M6 nuts. **h.** Detail of the Raspberry pi HQ camera with the lens assemble. **i.** Tightening the coupler with the lead screw after inserting it on both platforms.

### 5.10. Soldering

1. Glue the electronic box components together (Figure 17a).
2. Attach the ON/OFF switch and the 2.1 mm barrel jack on the electronics box (Figure 17a).
3. Cut at the right size and solder the pin male headers (Figure 17b).
4. Attach the two big (5W) potentiometers using the two holes of the PCB (Figure 17b).
5. Solder the pins of the 5W potentiometers using cables (Figure 17b).
6. Solder power cables between the barrel jack and the ON/OFF switch (Figure 17c).
7. Solder the four 10k (small) potentiometers on their respective places (Figure 17c).
8. Solder the Phoenix screw (green) connectors or pins on the connections for the IR LED cables (Figure 17c).
9. Solder the Vcc and GND wires and connect them with a power jack connector (Figure 17c).
10. You can attach the base of the Arduino Mega to a 10 mm PMMA or other plastic sheet to keep it flat (Figure 17c).
11. Solder 5 screw connectors on the 1-6 connector PCB and attach the PCB with a laser cut 1-6 connector base using four M2 screws and nuts (Figure 17e).
12. Solder the 82Ω resistors, power supply cables and IR LEDs on the matrix (Figure 17e).
13. Check if the LEDs are working. You can use your mobile camera to catch the infrared light (Figure 17e-f).
14. Attach the matrix to one of the arena sheets using two M3 screws (Figure 17e-f).

**Figure 17.**
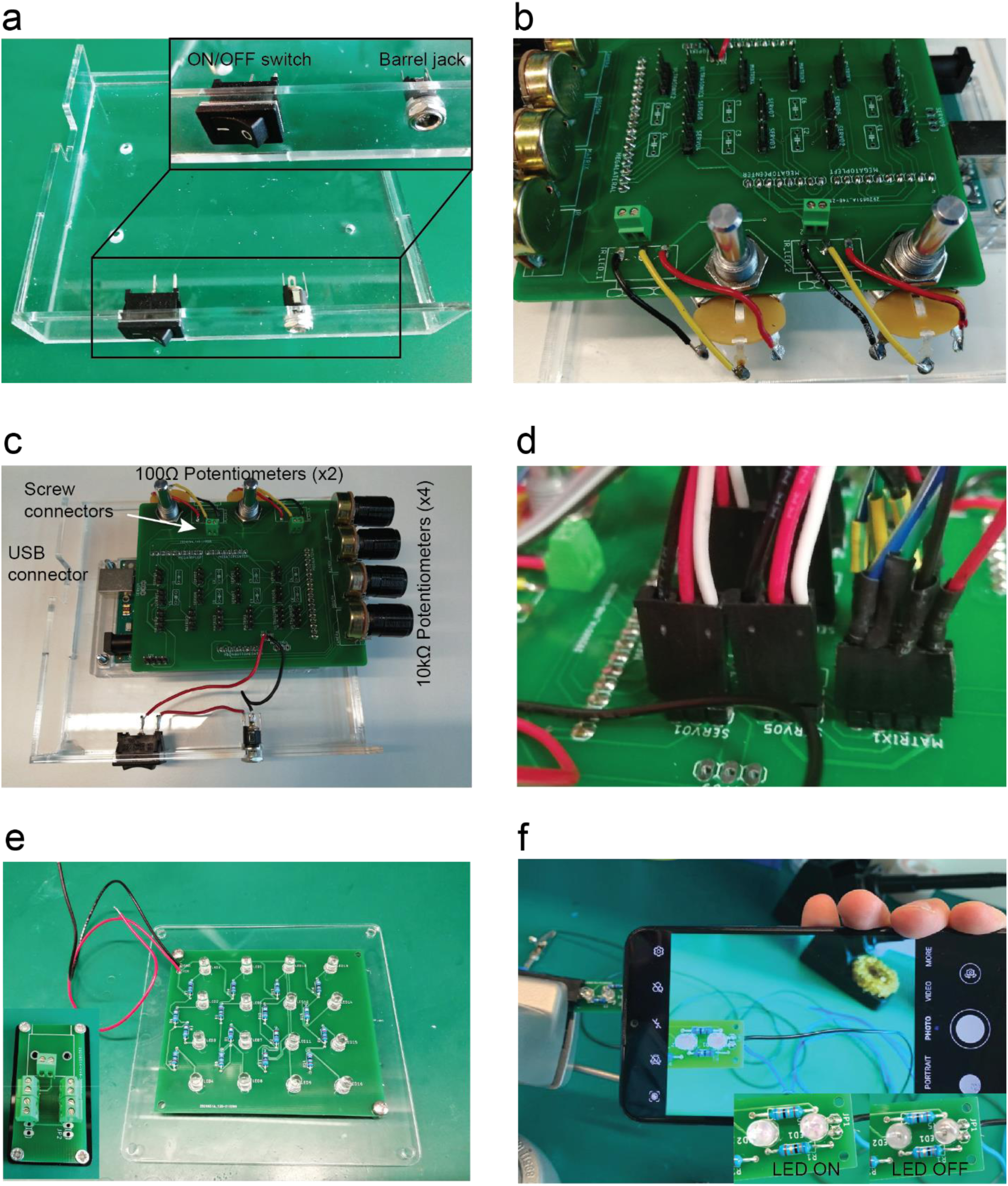
Control board shield soldering. **a.** Solder the four 10 kΩ potentiometers and attach the two 100 Ω potentiometers using the two holes located on the top of the PCB and the male pins. **b.** Solder the screw connectors to the potentiometers. **c.** Solder the pins of the two 100 Ω to the PCB pads. **d.** Detail of the cables connections and colour codes. **e.** Five screw connectors soldered on the 1-6 connector PCB and Matrix with soldered infrared LED and 68Ω resistors. **f.** The phone camera can be used to evaluate if the IR LED are on when we apply power (5V DC).

### 5.11. Electrical connections

The electrical connections are described in Figure 18. There are four possible connections to the RGB matrices but in the standard configuration we only connected two. The four IR linear arrays are connected to an intermediate PCB called 1-6 connector and from there to the 1-6 terminal connection. The two ultrasonic sensors are connected to the four pin connectors labelled “ultrasonic”. There are nine available connections to servo motors also labelled: “servo1-9” which are coincident with the number of servo objects on the Arduino program. In the standard configuration only seven are used, the other two servo motors are for an extra two matrices. The pi camera is connected to the raspberry pi using the ribbon cable. The Arduino is then connected to the raspberry pi or to a computer. Finally, connect the power supply to the PCB Jack connector. The raspberry pi must be connected to a keyboard and mouse using the USB ports and a computer display using the HDMI port.

**Figure 18.**
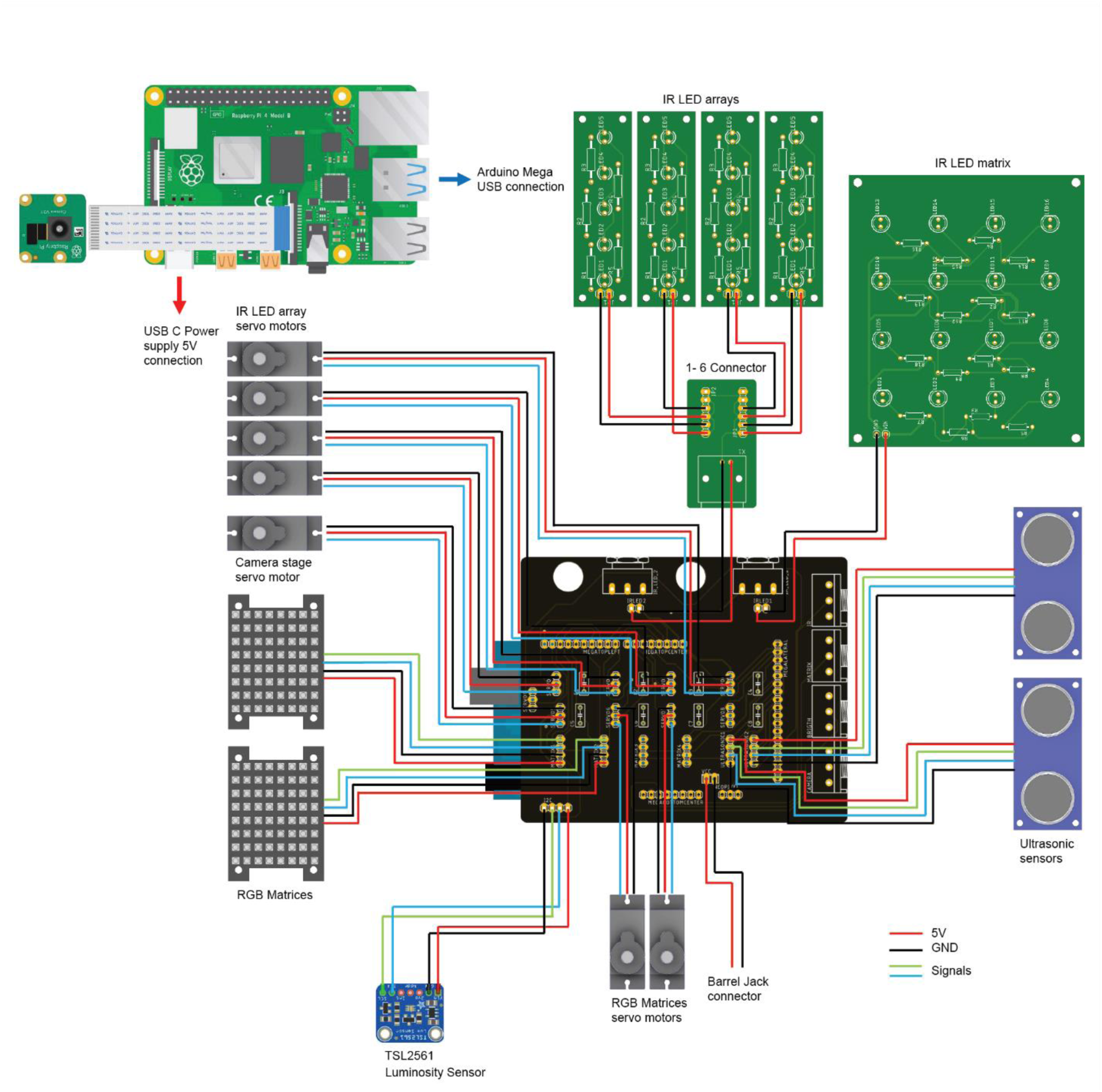
Electrical connections. The wiring required for all the actuators, sensors and illumination with the main control board.

Most of the power required to operate the device comes from the illumination. The current that the IR LED draw adding the four arrays is 1 A and increases up to 1.8A if you add the matrix illumination from below the arena. Every RGB matrix operating at maximum brightness demands between 1 – 1.2 A depending on the colour. The Dotstar matrices can go to 2.5 A combining the three RGB channels. A high demand of light intensity requires high currents, if that is the case it is important to supervise the device and make sure there are no exposed conductors.

## 6. Operation Instructions

This section has the following subsections:

6.1 Programming the Arduino Mega 2560 in Windows systems
6.2 Relationship of functions in the Arduino code
6.3 Acquiring video recordings from the Raspberry Pi
6.4 Analysing videos with BIO
6.5 Running BIO in on-line mode

### 6.1. Programming the Arduino Mega 2560 in Windows systems

1. Download and install Arduino environment on the computer (www.arduino.org).

– Install the Adafruit GFX library.
– Install the Adafruit DotStarMatrix library.
– Install the Adafruit DotStar library.
– Install the Adafruit sensor library.
– Install the Adafruit TSL2561_U library.
– Install the Adafruit AS7341 library.
– Install the elapsedmillis library.
– The other required libraries SPI, Wire (I2C) and servo are included in the standard libraries of the Arduino IDE.
2. Open the Arduino script:

– OptoPi.ino
3. From the ‘‘Tools” tab:

– Select from ‘‘Boards” the ‘‘Arduino Mega or Mega 2560”.
– Select from ‘‘Processor” the ‘‘ATmega2560”.
– Select from ‘‘Port” the ‘‘COM (Arduino Mega or Mega 2560)”.
4. Compile and upload the code (clicking on the sideways arrow).
5. The arena is ready to be used.

### 6.2. Relationship of functions in the Arduino code

The main functions can be divided in two main groups: setting functions and automated functions.

The setting functions enable the control of the device elements using the knobs and interfaces. This mode of operation is also used to read the values of of the sensors by opening the serial port (Tools -> Serial monitor). These functions can be set-up once and the device can then be used by just using the knobs. This is the mode of operation for simple recording experiments, or to find the right settings for a future automated experiment.

The automated functions are those that enable using the device with automatic experimental protocols that actuate the device without the user intervention. For example, it can create an optogenetic blinking series - *blink()* function - or an intensity ramp – *intensityramp()* function -. The parameters that receive the functions to operate are global variables declared at the top of the program. After the *setup()* function that is always executed once, on the main *loop()* function, the program waits for the user to press “space” key from the keyboard connected to the Raspberry pi. The values printed are read by the Raspberry Pi and saved into a txt. file with timestamps.

1. Setting functions. The setting functions are functions intended to be used to adjust parameters of the setup or simply control the setup manually without any protocol automation.

- *void* Settings () This function prints on the serial port the distances of the two ultrasonic sensors (Top and bottom), the camera distance, the angle of the two light sources (IR and RGB) and sets the RBG matrices brightness and at the same time is reading the knob values so the user can control all those parameters and adjust them.
- *void* read_irradiance () This function reads the value sent by the TSL2561 and converts the illuminance into irradiance using the calibration equations. Depending on which of the boolean variables red, green, blue are set to true the specific calibration equation will be choosen.
- *void* read_spectral_sensor () It reads the incoming data sent through the I^2^C bus and prints on the serial port the normalized value for each wavelength from 415nm to NIR (optional if connecting the Adafruit AS7431 module).
- *void* Set_Matrix_Color_Brightness (*byte* Red_int, *byte* Green_int, *byte* Blue_int, *byte* brightness) The function receives four input parameters of the byte types, the first three are variables that can be found at the beginning of the program, they correspond to the value from 0 to 255 of red, green and blue in that order. The fourth byte is the brightness also from 0 to 255.
2. Automated functions. The automated functions are required only if automatic optogenetic stimulation is required, If the user wants to control the setup manually, these functions are not required. These functions cannot be used at the same time as they control the RGB matrices in a different way.

- *void* intensity_ramp (*byte* max_value, *unsigned int* interval) The ramp function increases linearly the value of the light intensity up to a maximum value (max_value) given a certain interval which is defined at the beginning of the program as DELAY (Figure 19a-b).
- *void* blinking (*unsigned int* interval_low, *unsigned int* interval) The blinking function generates ON/OFF cycles with a fixed light intensity and an initial waiting period defined as DELAY_0 at the beginning of the program. In this case the variable DELAY is also used as interval between the ON and the OFF cycles (Figure 19c-d).

**Figure 19.**
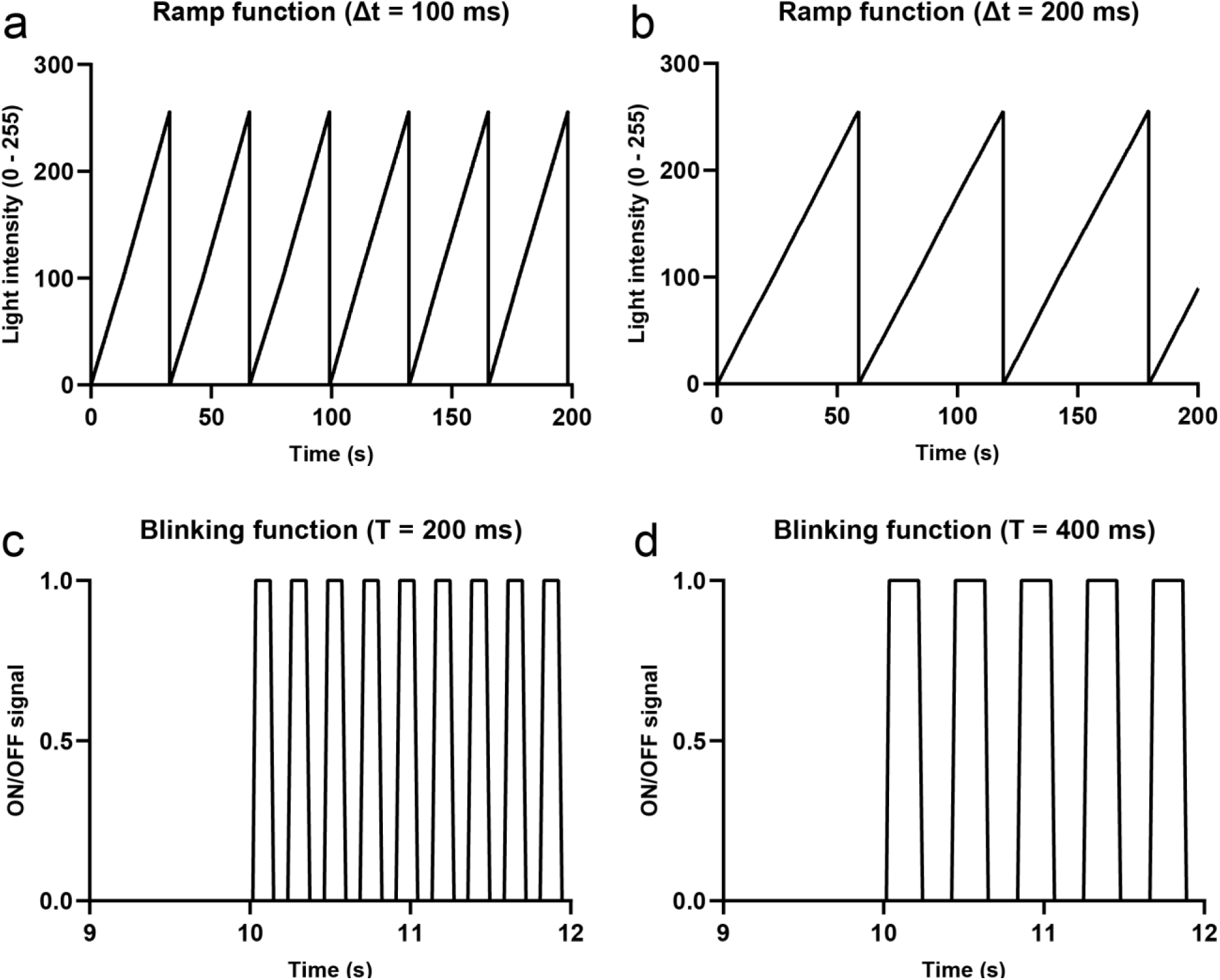
Automated functions. **a.** Ramp profile of light intensity, created by increments of one byte spaced 0.1 seconds. This pattern is generated by the automated function intensity_ramp(). **b.** The same function called with 0.2 seconds interval. **c.** ON/OFF cycles generated with both RGB matrices with a period of time of 0.1 seconds between ON and OFF states and an initial time of 10 seconds in off position. This pattern is generated by the automated function blinking. **d.** The blinking function using a delay of 0.2 seconds delay.

### 6.3. Acquiring video recordings from the Raspberry Pi

Videos in this manuscript have been acquired through two methods: 1) using the RaspberryPi 4 via a Python script (‘Camera_acquisition.py’) operating on a Raspbian OS, and 2) directly via BIO on an Ubuntu OS (see 6.5 below).

The ‘Camera_acquisition.py’ communicates with the Pi camera module to enable the user to modify video acquisition parameters including framerate, frame resolution and frame colour. As such, the ’Camera_acquisition.py’ script can be easily modified to meet a user’s needs. To simply record a video, lines 8, 21-28, 39, 41 and 45-51 can be commented out. However, should the Arduino’s ‘state’ need to be recorded in addition to a video, for instance, if an optogenetic stimulus is being delivered, keep the ‘Camera_acquisition.py’ script unedited. For synchronisation between the Arduino and the RaspberryPi first upload the Arduino script (‘OptoPi.ino’), commenting out the desired protocol, for instance, *settings()* or *blinking()*. Having ensured that the OptoPi.ino script has uploaded correctly (receiving a prompt from the Arduino IDE) simply run the ‘Camera_acquisition.py’ script via your chosen Python IDE. This will prompt the user with an unrecorded preview enabling the user to make any modifications before video acquisition begins. The user is prompted to press the ‘Enter’ key to begin recording and upon doing so the Raspberry Pi will send a serial command to the Arduino. As currently written, when the ‘OptoPi.ino’ is uploaded to the Arduino, the Arduino is essentially kept in an inactive state until prompted by a serial command. Therefore, upon pressing the ‘Enter’ key, the RaspberryPi sends this serial command thereby activating the Arduino to begin the selected protocol. The RaspberryPi will next create a start time point and begin recording the video. As a part of enacting its selected protocol, the Arduino will send via serial port a string detailing its current ‘state’, for example, LED matrix intensity. This information is decoded by the RaspberryPi and written to a .csv file along with a timestamp thereby providing information as a series of timestamps as to when stimuli are being delivered by the Arduino. This process is repeated with the RaspberryPi assessing whether the time elapsed exceeds a user defined experiment duration (the variable ‘experiment_duration’). Once the required time has elapsed the RaspberryPi ceases recording the video. As a part of debugging, a user may need to upload their ‘OptoPi.ino’ several times prior to running the ‘Camera_acquisition.py’ script to ensure that the Arduino is in an ‘inactive’ state and awaiting the appropriate serial command from the RaspberryPi.

The steps to take are the following:

1. Install the required python libraries, these are as follows:

– Serial
– Picamera
– Time
– System
– Datetime

The only one you will probably have to install is the serial library. But first you must ensure you have python3 installed by typing the following command in the terminal: *python 3 – version*

Should this return a value, that means you have correctly installed a version of python3 and may proceed. Otherwise, type the following commands:

*sudo apt-get install python3.8 python3-pip*
*sudo apt install python3-pip*
Then you install the pySerial library by typing
*python3 -m pip install pyserial*

2. Open the terminal and write: *ls /dev/tty\** and press enter. This will open the list serial ports available. If you do that before and after connecting the Arduino board, the name that only appears when the Arduino is plugged is the name of the Arduino serial port usually ‘/dev/ttyACM0’, ‘/dev/ttyUSB0’, or similar. Alternatively, copy the name of the port the Arduino is connected to via the Arduino IDE by clicking the dropdown ‘Tools/Ports’ and find your connected Arduino.

3. Copy the “Camera_aquisition.py” inside a folder on the Raspberry Pi desktop and open the script in “Camera_aquisition.py” via Thonny, a Python IDE already available on Raspbian, or any other IDE of your choice.

4. In the script modify the “Acquisition_time” and write a “file_name” of choice and press RUN. A window with the video recording will open and a .CSV file with the name you assigned will be automatically created and the timestamps of data sent from the Arduino code will be logged in for the duration of the acquisition.

### 6.4. Analysing videos with BIO

Once the video has been acquired it can be analysed using multiple software. Here we present BIO, which enables tracking multiple individuals simultaneously, while keeping individual’s identity upon collisions. BIO has been tested and works across operating systems including Windows and Ubuntu (20.04 and Ubuntu Desktop 21.10 on Raspberry Pi 4). For Windows systems, BIO can be downloaded and installed from https://joostdefolter.info/bio-research. For Ubuntu, BIO can be built following the instructions outlined in https://github.com/folterj/BioImageOperation/blob/master/qt%20opencv%20linux%20installation%20notes.txt. To run BIO on Windows simply double click its executable. To launch BIO on Ubuntu, first open a terminal session and navigate to the cloned GitHub directory (likely still named BioImageOperation). From here, navigate to the build/BioImageOperation directory. Finally, launch BIO via the command : *./BioImageOperation*.

A description of the steps required to generate a tracking script and the functions available within BIO are available at https://joostdefolter.info/bio-research. Additionally, the dropdown Help -> Script Help points towards different functions available to the BIO user. As a starting point, BIO generates a template script by clicking the dropdown ‘Script -> Generate tracking script’ and subsequently selecting a video to be analysed. Once this has been generated the user must follow the steps laid out in this script to first create a background image, assign appropriate thresholds to properly track the objects of interest and finally output these tracks as data output and/or video. Additionally, the user can run BIO when it includes the ‘Debug()’ function, this will output estimates for Threshold(), CreateClusters() and CreateTracks() parameters. These serve as useful starting points and aid in the user’s finding an optimal set of values for each parameter. Throughout optimisation of tracking parameters the function ShowImage() should be included in the script in order for the user to receive visual feedback.

Example scripts used to track videos are provided, but a brief description of how they were tracked is as follows. Videos are read into BIO via the *‘OpenVideo()*’ command which accepts a string detailing the path to the chosen video. This argument also accepts an ‘*interval=n’* flag wherein the BIO will only use every nth frame. For videos acquired at higher framerates or where there is very little difference between successive frames, including this interval will speed up analysis. BIO currently analyses 8-bit videos, therefore it is necessary to convert the video using the *Grayscale()* argument. Following this, *AddSeries()* and *GetSeriesMedian()* are used to generate an ‘average’ background image. Provided all the potential subjects of interest moved during the video you will be left with a background only including the static objects present in the video. Save this background image. BIO accepts multiple image formats, therefore we recommend the use of a lossless compression image format, such as .png and .tiff. Assign the background image with an appropriate variable name, e.g. ‘*background = OpenImage(‘filepath.png’)*’. It is this background that potential objects need to be compared against using the ‘*DifferenceAbs(background)*’ function. The precise threshold, which is a percentage that any object of interest must differ to the background, is user defined and is a float from 0 to 1. The higher the contrast, the more stringent the threshold (i.e. higher) can be. For the example videos provided we used thresholds below 0.1.

At this stage a mask can also be provided. Essentially a mask dictates to BIO over which pixels it should attempt to track. This mask must be an 8-bit binary image of the same dimensions as the background image. Pixels with a 255 value will be the pixels searched in the video when tracking, while those with a value of 0 will be ignored. Such binary images are easily made by programs such as Inkscape (https://inkscape.org/) and FIJI (https://imagej.net/software/fiji/). Care must be taken to ensure the image is 8-bit. The mask can be read into BIO via the following command: ‘*mask = OpenImage(‘mask.png’)*’. In order to use the mask, use the function *Mask(mask)* after the *Threshold()* function.

The final steps involve assigning clusters, to delineate which objects to track, and the actual tracking. Assign a minimum and maximum value to the *CreateClusters()* function e.g. *CreateClusters(MinArea=80,MaxArea=1000)* for BIO to detect objects that reach the contrast threshold and that consist of between 80-1000 pixels (in this example). Finally, assign values to the *CreateTracks()* function to determine the maximum acceptable movement of a cluster per frame, the minimum active number of frames a cluster should be moving before it is tracked and finally the maximum number of frames a cluster is inactive before it is no longer tracked. For example, *CreateTracks(MaxMove=20,MinActive=3,MaxInactive=3)*. The movement speed of the organism being tracked is a critical determiner of what *MaxMove* value should be with faster animals requiring a higher *MaxMove* value. Once these settings have been optimised it is merely a matter of saving the track outputs. Should the user simply want the data use the function ‘*SaveTracks()*’ and write the data output to a csv file. However, if the user additionally wants to save the tracking as a video use the functions *DrawTracks()*, changing the flags to their specification, and finally output a video with the *SaveVideo()* function.

Note, that using a wide-eye lens to record animals in a wide arena will produce videos with barrel distortion, i.e. objects and distances in the centre of the arena appear larger than those towards the edges. For the experiments performed for this publication arena sizes were not large enough for such barrel distortions to become an issue. However, for accurate tracking in wider arenas it is essential to correct these distortions before tracking the videos in BIO. Such corrections can be performed using BIO by first imaging a checkerboard pattern under the camera position conditions used for experiments and performing the OpticalCalibration() function on images and videos. Example checkerboard image with distortion, with distortion corrected and example scripts are available in Supplementary folder visual_distortion_correction.

### 6.5. Running BIO in on-line mode

The steps required to run BIO online are the same as outlined above, but rather than selecting a single video, the user must use ‘*OpenCapture(0)*’. If BIO detects a functioning camera, it will read a live feed from it. When using the on-line mode, one way to compute a background image is to run BIO and have it iteratively generate a median frame over a suitable period of time (*example_script.bioscript*). Save this background image and proceed to run BIO as one would during offline analysis. Additionally, BIO comes with an *UpdateBackground()* function, using this the user can take the first frame as a background and proceed to update it as the experiment proceeds.

## 7. Validation and Characterization

This section has the following subsections:

7.1 Experimental characterisation of the illumination and recording modules
7.2 Technical characterisation of the optogenetic modules
7.3 Technical characterisations of the precision of the device
7.4 Experimental characterisation of the optogenetic modules

### 7.1 Experimental characterisation of the illumination and recording modules

To test the OptoPi’s performance, we utilised various lighting settings to record and analyse the behaviour of *Drosophila melanogaster* adult flies and 1^st^ instar larvae, as well as *Danio rerio* fish larvae (Figure 20). We recorded animals in ‘ambient’ lighting conditions, wherein the overhead LED matrices emitted white light, and infrared lighting, either through darkfield illumination from the side IR LED matrices or backlighting from IR LEDs beneath the stage. We found that ambient lighting worked particularly well for adult flies and larval fish, as their dark pigmentation provided good contrast against the dark background. However, the small and transparent body of 1^st^ instar fly larvae did not provide enough contrast under ambient light conditions, and these are best recorded using darkfield illumination from the side IR LEDs.

**Figure 20.**
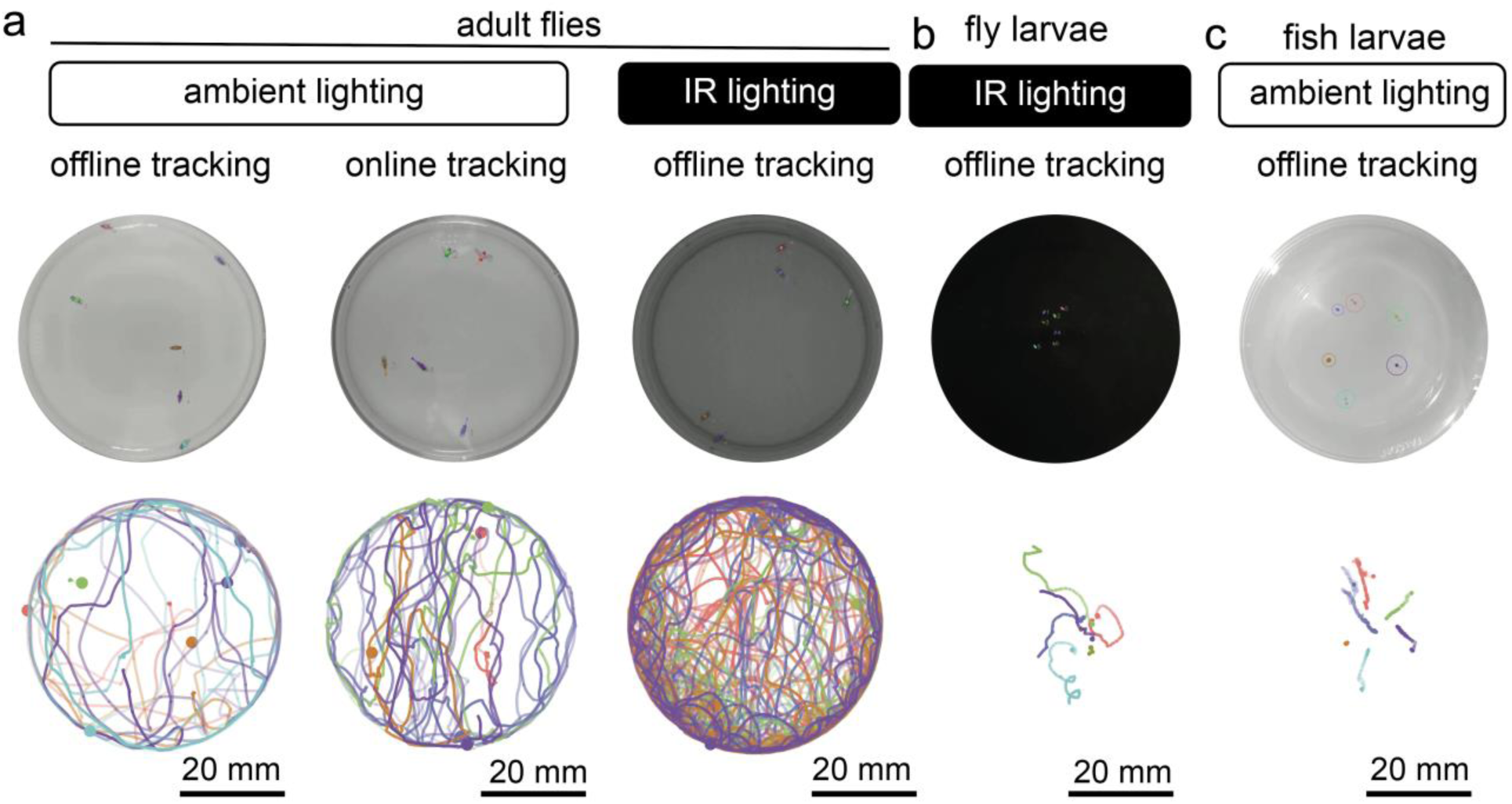
Experimental characterisation of the illumination and recording modules. **a.** Example *D. melanogaster* adult snapshots (upper panels) and trajectories (lower panels) under different lighting and recording conditions (left and middle under ambient lighting, right in infrared lighting, middle column is tracked live). **b**. Example *D. melanogaster* larvae free-roaming in infrared lighting, example image (upper panel), total trajectories (lower panel). **c.** Example free-roaming *D. rerio* larvae in ambient lighting showing example image (upper panel) and trajectories (lower panel).

To demonstrate that each of these conditions enables animal tracking we analysed these experiments using BIO both online and offline (Figure 20 and Supplementary Videos 17-24). Analysis and visualisation of larval trajectories were done in R relying on the package trajr (https://cran.r-project.org/web/packages/trajr/index.html).

### 7.2 Technical characterisation of the optogenetic modules

To characterize the capabilities of the optogenetic modules, the irradiance and illuminances were simultaneously measured on the set-up using a calibration platform with the in-built TSL2561 illuminance sensor and a calibrated Thorlabs S120VC photodiode placed on the centre of the arena (Figure 21). The Thorlabs S120VC was connected to a PM100D compact power meter console. The position of the two RGB matrices were at an angle of 100°, Bottom platform distance (2 cm) and distance of the top platform (11 cm). That position was selected as the maximum stimulation achievable with two RGB Adafruit DotStar high density matrices. The maximum irradiances measured are: 4.22, 5.88, 5.84 µW/mm^2^ for red, green and blue respectively using two RGB high density grids. Then both functions (irradiance and illuminance) were fit in quadratic equations (R^2^ = 0.9992 - 0.9999) and the two functions were used to generate values to correlate the measurements of both sensors (Figure 20b) with also high correlation coefficients (R^2^ = 0.9994 - 1). These equations have been introduced to the Arduino code inside the *read_irradiance()* function and can be used by the user to find the adequate position for the light stimulation. Depending on how the initial boolean variables for red, green, or blue are set, the function will convert the TSL2561 reading using the corresponding equation. The measured averaged minimum change in irradiance in each channel is 0.024, 0.033, 0.033 µW/mm^2^ for the LEDs red, green and blue respectively. The residual irradiance (Bright byte value 0) was measured as 0 1.5·10^-4^ µW/mm^2^.

**Figure 21.**
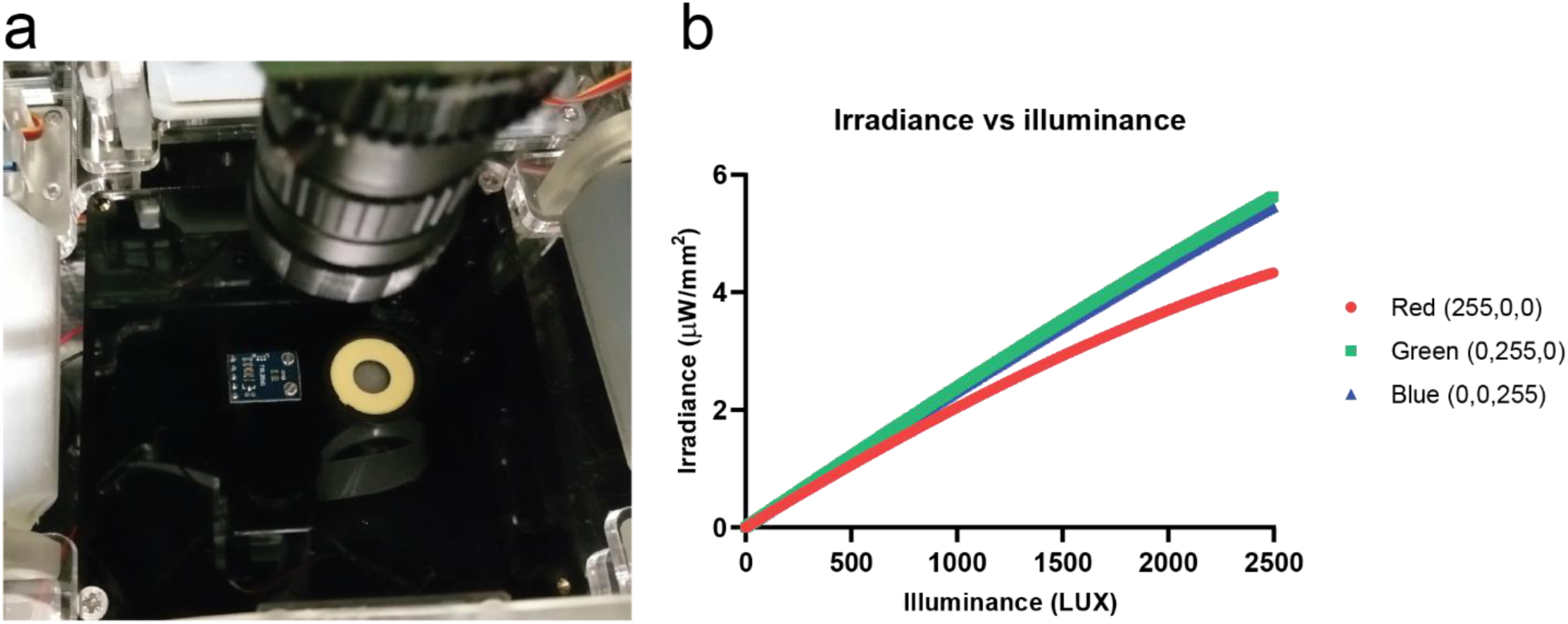
Technical characterisation of the optogenetic module. **a**. Irradiance characterization setup, the TSL2561 luminosity sensor (left) and the Thorlabs S120VC (Right) on the calibration arena. **b.** The calibration functions used to find the linear regressions for the red (609 nm), green (526 nm) and blue (455 nm) lights.

### 7.3 Technical characterisations of the precision of the device

The following table collects the main features of the device regarding the resolution of the different systems that measure the angular, horizontal, and vertical positions of the light sources as well as the camera. The maximum irradiance measured for red, green and blue light and their resolution, and the measured wavelengths for the optogenetic stimulation light sources and for the infrared illumination.

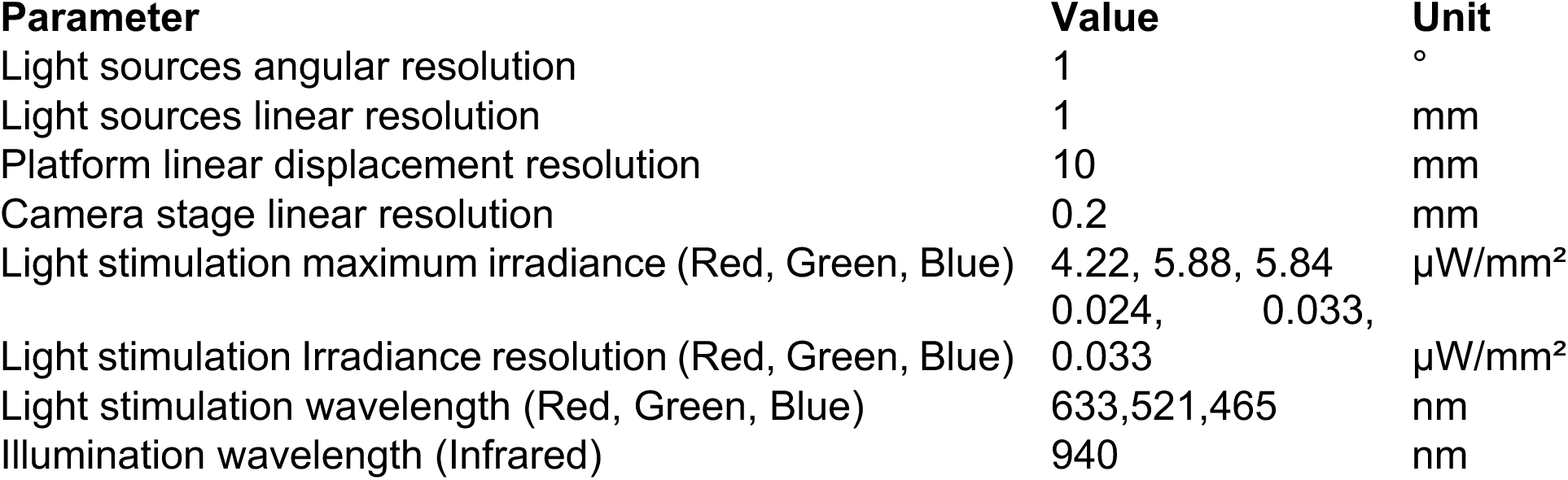

### 7.4 Experimental characterisation of the optogenetic modules

Here we validate experimentally the optogenetic capabilities of the OptoPi by using two stereotypical behaviours of the *Drosophila melanogaster* larvae: rolling and stopping.

Rolling is a nocifensive behaviour displayed by larvae when their nociceptive cells are being stimulated either mechanically or thermally. It is possible to elicit this behaviour without the stimuli that naturally trigger it by optogenetically activating either the sensory neurons, or downstream neurons of the nociceptive circuit. Here, we employed a driver line, *GMR72F11-Gal4,* which is expressed in interneurons of the nociceptive circuit and whose optogenetic activation has been shown to elicit robust rolling behaviour[11]. To activate these neurons we crossed this Gal4 to a UAS line encoding the red light activated ion channel CsChrimson (*UAS-CsChrimson*, BL79599)[7].

Stopping is a natural behaviour that larvae perform when they encounter an aversive stimulus, or during regular navigation when re-evaluating their direction of travel. However, stopping can also be elicited artificially by simultaneously inactivating all motorneurons, thus rendering the animals immobile. We employed this later approach by targeting all motorneurons (and a few other interneurons) using the vGlut-Gal4 (BL26160) driver line [12], [13]. We crossed this line to UAS-GtACR1, which encodes a green-light sensitive chloride channel, which upon activation by light inhibits the neurons where is being expressed[12].

For either experiment, 30 male and female adults were placed in cages placed on top of a grape juice agarose plate (3% agar, 25% grape juice) for 4 hours. The plates were smeared with yeast paste containing 1mM all-trans retinol (Sigma Aldrich, CAS116-31-4). Larvae for optogenetic experiments were collected 48 hours following adult introduction to cages. *Drosophila* crosses were reared at 25C and in the dark. The experimental larvae were collected from the grape juice agarose plates using forceps, cleaned with water, and placed over a fresh agarose plate (1%). During behavioural assays, larvae were kept in the dark and intermittently exposed to red/green light via the blinking (20 seconds on 20 seconds off) protocol in OptoPi.ino. Thus, during periods when the larvae were in the dark they could behave normally while exposure to red/green light optogentically activated target neurons. *GMR72F11-Gal4/UAS-CsChrimson* larvae exhibited rolling behaviour only during red light stimulation and *VGlut-Gal4/UAS-GtACR1* larvae paused in response to green light stimulation (Figure 1, Table 1, Supplementary Videos 22-23). Periods of exposure to red/green light are exported into a separate .csv file or annotated directly on the video. These behaviours can also be easily observed by colour coding their tracks dependent on whether they were being exposed to red/green light or in the dark (Figure 22).

**Figure 22.**
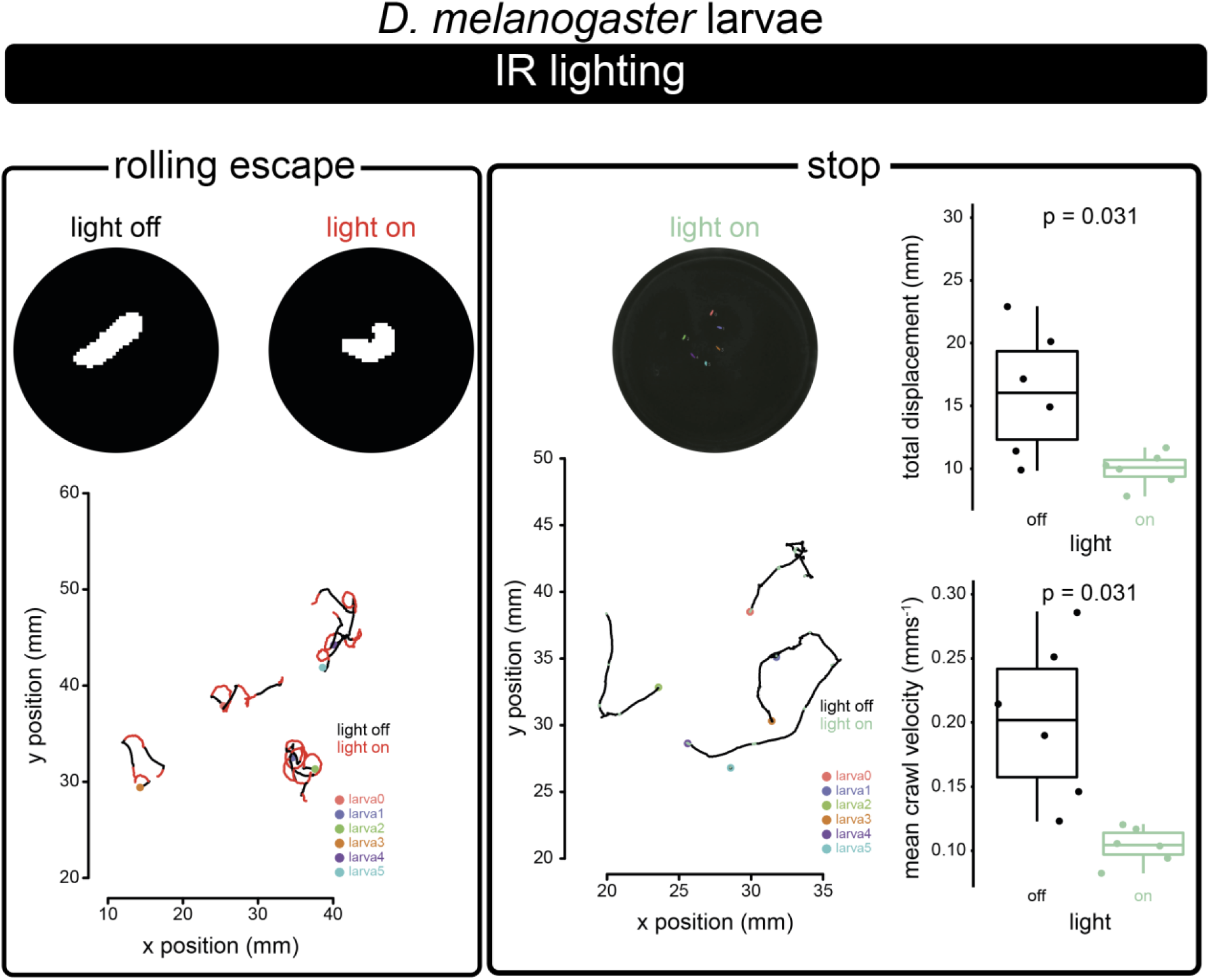
Experimental characterisation of the optogenetic module. Example *D. melanogaster* larvae optogenetic experiments. Left: Rolling escape behaviour elicited by exposing *GMR72F11-Gal4/UAS-CsChrimson* larvae to red light. Red portions of trajectory occur during exposure to red light. Upper panel left shows larval shape without red light, upper panel right shows larval shape with red light. Right: Stopping behaviour elicited by exposing *VGlut-Gal4/UAS-GtACR1* larvae to pulses of green light. Significance determined by paired Wilcoxon test performed using R package ggpubr. Lower left, green portions of trajectory occur during exposure to green light. Upper right panel, total larval displacement during periods of time when green light is off vs on. Lower right panel, mean crawl velocity during periods of time when green light is off vs on.

**Table 1.** Example settings for OptoPi that can be used to reproduce the example experiments of the figure. These combine settings determined by OptoPi.ino and user defined settings. Numbers in grey refer to values that are inessential to the running of the relevant experiment e.g. IR experiments without RGB lighting.

| Experiment | Db | Dt | IR | RGB | Cd | IR Br | IR Lens | RGB Lighting | RGB Br | RGB lighting protocol |
| --- | --- | --- | --- | --- | --- | --- | --- | --- | --- | --- |
| A – *D. mel* adult offline ambient | 2 | 7 | 45 | 180 | 138 | 100 | off | 255,255,255 | 80 | constant |
| A – *D. mel* adult online ambient | 2 | 7 | 45 | 180 | 138 | 100 | off | 255, 255, 255 | 80 | constant |
| *D. mel* adult offline IR | 1.4 | 3 | 66 | 142 | 69 | 100 |  |  |  |  |
| *D. mel* 1<sup>st</sup> instar larvae IR | 3.8 | 7 | 98 | 180 | 0 | 100 | on | off | 0 | Constant |
| *D. rerio* offline ambient | 4 | 7 | 98 | 180 | 0 | 100 | off | 255, 255, 255 | 80 | constant |
| *D. mel* larvae GMR72F11/C sChrimson IR | 4 | 7 | 98 | 180 | 70 | 100 | on | 255,0,0 | 80 | blinking (20s on, 20s off) |
| *D. mel* larvae VGlut/GtACR 1 IR | 3.8 | 7 | 98 | 180 | 70 | 100 | on | 82, 255, 0 | 80 | blinking (20s on, 20s off) |

We therefore demonstrate that the OptoPi reliably elicits well-characterised optogenetically induced behaviours.

## 8. Acknowledgements

We are thankful to Alexander Borst for sharing their GtACR1 flies and Rashmi Priya for providing the zebrafish larvae. L.L.P.-G.’s laboratory was supported by a European Research Council (ERC) Starting Investigator Grant (802531), a Human Frontiers Science Grant and an Allen Institute Distinguished Investigator Award. RJVR is supported by a Boehringer Ingelheim Fonds PhD fellowship. Work in the Francis Crick Institute is supported by core funding from Cancer Research UK (FC001594), the UK Medical Research Council (FC001594), and the Wellcome Trust (FC001594). For the purpose of Open Access, the authors have applied a CC BY public copyright license to any Author Accepted Manuscript version arising from this submission.

## 9. Declaration of interest

The authors declare that they have no known competing financial interests or personal relationships that could have appeared to influence the work reported in this paper.

## 10. Human and animal rights

All procedures were performed in accordance with the UK Animals (Scientific Procedures) act 1986 and approved by the animal welfare committee of The Francis Crick Institute.

## Notes

https://github.com/PrietoGodinoLab/OptoPi

https://osf.io/pu8f5/

### Competing Interest Statement

The authors have declared no competing interest.

### Summary of Updates

I realised that the pdf I posted had lost the links to the supplementary videos. This new version has the links intact. The rest of the manuscript is identical.

